# Endocrine–Neural Interactions Regulate Antral CCK2R⁺ Stem Cells in Gastric Inflammation and Preneoplasia

**DOI:** 10.64898/2026.08.28.747959

**Authors:** Biyun Zheng, Hualong Zheng, Ruhong Tu, Fenglin Chen, Jiaoqian Lu, Hiroki Kobayashi, Puran Zhang, Yi Zeng, Guodong Lian, Feijing Wu, Xiaozhong Wang, Xiaofei Zhi, Kexuan Huang, Jin Qian, Quin T. Waterbury, Shuang Li, Juli Lin, Xinliang Xiong, Ermanno Malagola, Yosuke Ochiai, Masahiro Hata, Junya Arai, Leah B. Zamechek, Timothy C. Wang

## Abstract

Antral CCK2R⁺ stem cells are regulated by gastrin, but how endocrine and neural cues integrate under chronic injury remains unclear. Here we show that inducible hypogastrinemia shifts from asymmetric renewal to symmetric expansion of CCK2R⁺ stem cells. With carcinogenic stress, these cells acquire a cycling, injury-responsive progenitor state revealed by single-cell RNA profiling. Acute gastrin loss activates a CCK2R⁺ nodose–DMV vagal reflex that increases acetylcholine release, NGF production, cholinergic innervation, and Chrm3 expression, driving ERK and YAP signaling in CCK2R⁺ stem cells. Vagotomy, Trk inhibition, or Chrm3 deletion each suppressed stem-cell expansion. In H. pylori and MNU injury models, hypogastrinemia amplified inflammation, dysplasia, and CCK2R⁺ clone expansion, whereas gastrin suppressed these responses. Human scRNA-seq and spatial profiling confirmed G-cell depletion and progenitor-state enrichment. These findings define an endocrine–neural–epithelial axis in which gastrin loss boosts vagal–M3R signaling to initiate antral preneoplasia, highlighting this pathway for early interception.

**In brief:** Zheng and colleagues show that gastrin loss is sensed by CCK2R⁺ vagal afferents to engage a nodose–brainstem cholinergic reflex that activates epithelial M3R–ERK/YAP signaling, while lifting gastrin’s cell-intrinsic brake on symmetric division. Together these inputs expand antral CCK2R⁺ +4 stem cells, drive injury-state reprogramming, and promote preneoplastic progression.

**Highlights:**

- CCK2R⁺ vagal afferents sense gastrin loss and engage a nodose–brainstem reflex.
- Gastrin deficiency induces NGF–TrkA–dependent cholinergic remodeling and elevates epithelial M3R.
- Cholinergic M3R signaling activates ERK and YAP to drive expansion of antral CCK2R⁺ +4 stem cells.
- Direct loss of gastrin’s restraint on symmetric division synergizes with indirect neural signaling to promote preneoplasia.

## INTRODUCTION

The distal antrum is a major site of chronic inflammation and early gastric carcinogenesis, where a narrow isthmus houses stem and progenitor cells that drive bidirectional epithelial renewal.^1^ Under homeostasis, these stem cells divide predominantly asymmetrically, but chronic inflammation or genotoxic stress can shift this equilibrium toward symmetric expansion, promoting clonal amplification, metaplasia, and dysplasia.^2,3^ Yet how endocrine and neural pathways converge to regulate antral stem-cell behavior during chronic injury remains poorly defined.

Several useful markers for long-lived antral stem/progenitor populations have been defined in the murine stomach, including Lgr5⁺,^4^ SOX2⁺ and SOX9⁺,^2^ Axin2⁺,^5^ Bhlha15/Mist1⁺,^6^ AQP5⁺,^7^ Runx1 enhancer–marked,^3^ and CCK2R⁺.^8^ Among these, CCK2R marks a +4 isthmal stem-cell population that is largely distinct from Lgr5^hi^ basal cells and exhibits quiescence, label retention, and asymmetric renewal under baseline conditions.^8^ Under carcinogenic stress, however, CCK2R⁺ cells undergo symmetric expansion and can serve as putative cells of origin for antral tumorigenesis.^9^ Gastrin produced by adjacent G cells has been implicated in constraining this symmetric switch, but whether G-cell loss in vivo reprograms the niche to alter CCK2R⁺ lineage behavior has not been resolved.

The antral stem-cell niche incorporates endocrine, neural, epithelial, stromal, and immune cues.^1,10^ Gastrin directly restricts epithelial proliferation and biases CCK2R⁺ stem cells toward asymmetric renewal,^9^ but endocrine signals operate within a highly innervated environment. Vagal sensory afferents in the nodose ganglion (NG) directly detect gut-derived endocrine peptides—including GLP-1 from enteroendocrine L cells,^11^ CCK,^12^ and ghrelin^13^—which they relay to the nucleus tractus solitarius (NTS). The NTS then activates the dorsal motor nucleus of the vagus (DMV), forming a classical NG→NTS→DMV sensory–motor reflex that governs vagal cholinergic output to the gut.^14^ Immediately downstream of this reflex, muscarinic signaling—particularly through the M3 receptor (M3R)—is a key effector of vagal cholinergic output in the stomach. Previous work demonstrated that cholinergic nerves promote gastric epithelial proliferation and tumor growth through M3R activation, and that pharmacologic blockade of muscarinic signaling suppresses tumor initiation.^15^ However, this study did not define which epithelial populations receive M3R input, whether antral stem cells themselves express M3R, or how muscarinic signaling interfaces with G-cell loss, antral atrophy, or division-mode switching—leaving unresolved how cholinergic pathways intersect with endocrine regulation within the antral niche.

Neural regulation of stem cells is emerging as a conserved paradigm across tissues. CGRP⁺ nociceptive neurons cooperate with sympathetic fibers to maintain and mobilize hematopoietic stem cells,^16^ sympathetic overactivation depletes melanocyte stem cells and accelerates tissue aging,^17^ and enteric cholinergic neurons promote epithelial proliferation via muscarinic receptors.^15,18^ In addition, chronic stress–induced activation of the central amygdala-DMV pathway impairs intestinal stem-cell stemness and promotes premature aging, demonstrating direct brainstem control of gastrointestinal stem-cell behavior.^19^ In the stomach, vagal sensory and cholinergic pathways cooperate with gastrin to regulate corpus CCK2R⁺ progenitors during ulcer repair.^20^ Whether similar neuro–endocrine interactions operate in the G-cell rich antrum—and whether endocrine cues gate vagal circuits to regulate CCK2R⁺ stem cells during inflammation and neoplastic initiation—remains completely unknown. Here, we demonstrate that gastrin loss drives CCK2R⁺ stem-cell expansion through a dual mechanism: direct promotion of symmetric division and activation of a CCK2R⁺ vagal reflex that boosts cholinergic–M3R–ERK/YAP signaling. This defines an endocrine–neural–epithelial axis that primes the antrum for dysplasia and early tumor initiation.

## RESULTS

### G-Cell Loss Drives Expansion and Reprogramming of Antral CCK2R⁺ Stem Cells

To investigate how gastrin controls antral stem-cell dynamics, we crossed Cck2r-CreERT2; Rosa26-ZsGreen mice^8^ with Gastrin-DTR-p2A-tdTomato mice^20^, enabling inducible, selective ablation of mature G cells in adult animals—a key distinction from constitutive whole body Gastrin⁻/⁻ models.^9^ Diphtheria toxin (DT) efficiently eliminates G cells and induced sustained hypogastrinemia, whereas continuous gastrin infusion produces physiological hypergastrinemia.^20^ G-cell ablation caused a rapid and persistent expansion of antral CCK2R⁺ lineage-traced clones, detectable within 24 hours of induction and maintained for up to 9 months, while gastrin infusion markedly suppressed tracing (Figure 1A–C). Flow cytometry corroborated these changes (Figure 1D and S1A). CCK2R expression was largely restricted to the Lgr5^lo^ fraction^4^ of antral epithelial cells (Figure S1B–D), confirming that CCK2R⁺ stem cells correspond to the +4/upper-gland compartment rather than Lgr5^hi^ antral basal cells. In addition, CCK2R^+^ stem cells show higher levels of expression of other antral stem cell markers such as Sox2, Sox9 and Axin2 (Figure S1E). NUMB polarity analysis^21^ showed that G-cell loss shifted CCK2R⁺ stem cells toward symmetric divisions, whereas gastrin promoted asymmetric renewal (Figure 1E and S1F). Functionally, hypogastrinemia enhanced organoid growth, while gastrin suppressed organoid-forming capacity (Figure 1F and S1G-H).

**Figure 1.**
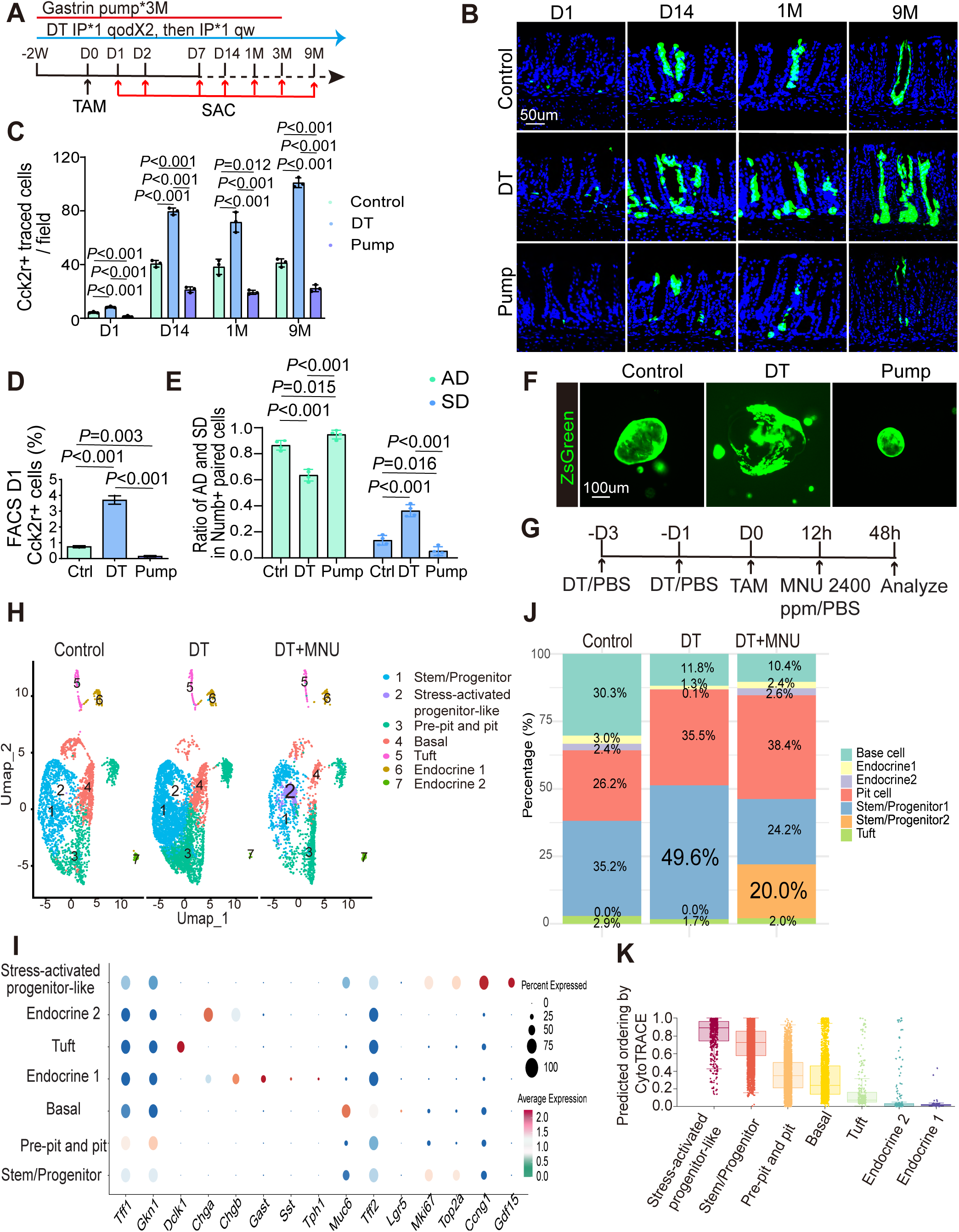
Hypogastrinemia reshapes antral CCK2R⁺ stem-cell behavior and transcriptional state toward stress and dysplastic reprogramming. (A–C) Inducible lineage-tracing strategy (A), representative antral images (B), and quantification of traced Cck2r⁺ cells at indicated time points after tamoxifen (C). (D) Flow cytometric quantification of antral Cck2r⁺ epithelial cells 1 day after tamoxifen. (E) Quantification of asymmetric (AD) versus symmetric (SD) division in paired ZsGreen⁺ cells 2 days after tamoxifen. (F) Representative organoids derived from FACS-sorted Cck2r⁺ cells after 7 days in culture. (G–I) scRNA-seq workflow (G) and UMAP visualization (H) of ZsGreen⁺ cells 2 days after tamoxifen, with marker-gene expression shown by dot plot (I). (J) Cluster proportions. (K) CytoTRACE scores across clusters. All experiments were performed in Cck2r-CreERT2; Rosa26-ZsGreen; Gastrin-DTR-p2A-TdTomato mice. Data are mean ± SD (n ≥ 3). Scale bars: 50 µm (B) and 100 µm (F). DT, diphtheria toxin; MNU, N-nitroso-N-methylurea; Pump, continuous gastrin infusion.

To define transcriptional consequences of gastrin deficiency, we performed scRNA-seq on FACS-sorted CCK2R⁺ cells at 48 h after tamoxifen induction from Cck2r-CreERT2; Rosa26-ZsGreen; Gastrin-DTR-p2A-tdTomato mice under three conditions: control (PBS), G-cell ablation (DT), and G-cell ablation with MNU exposure (DT+MNU) (Figure 1G). Unsupervised clustering of high-quality epithelial cells resolved seven transcriptionally distinct populations. Cluster identities were assigned based on canonical lineage markers and the top 50 differentially expressed genes per cluster (Figure 1H and 1I; Table S1). These included a stem/progenitor cluster (*Mki67, Ccna2, Top2a, Birc5, Stmn1,* Cluster 1); a stress-activated progenitor-like cluster (Cluster 2) enriched for p53 and stress-response genes such as *Gdf15*, *Trp53inp1*, and *Ccng1*, along with *Notch3* and *Mybl1*, which are associated with progenitor maintenance during regeneration; pre-pit and pit cell clusters (*Tff1, Tff2, Muc5ac*, Cluster 3); basal gland mucous cells (*Muc6, Lgr5, Aqp5, Glp1r, Car9,* Cluster 4); a tuft cell cluster (*Dclk1, Pou2f3,* Cluster 5); and two endocrine clusters—endocrine 1 (*Neurod1, Gast*, Cluster 6) and endocrine 2 (*Chga, Sstr2,* Cluster 7). Notably, the stress-activated progenitor-like cluster (Cluster 2) was present only in mice that had received DT+MNU and displayed elevated Mki67 and Top2a expression. Thus, these progenitor cells exhibiting elevated proliferative potential arose under severe regenerative or injury-induced conditions (Figure 1H). While G-cell ablation expanded the stem/progenitor cluster, DT+MNU uniquely induced the stress-activated progenitor-like cluster with the highest CytoTRACE scores and earliest Pseudotime position, suggesting that it represents a primitive progenitor state emerging under sustained injury stress (Figure 1J and 1K; Figure S1I-M), conditions shown to be associated with carcinogenesis.

Gene set enrichment analysis (GSEA) revealed that G-cell ablation alone activated interferon-α and interferon-γ response pathways, consistent with chronic inflammatory stress, while wild-type mice showed no histological changes for 9 months.^20^ In contrast, DT+MNU exposure further upregulated oncogenic and dysplasia-associated programs, including P53 activation, TNFα/NF-κB, and KRAS signaling (Figure S1N and S1O), indicating that carcinogenic stress synergizes with gastrin loss to drive transcriptional reprogramming toward a premalignant state. These findings demonstrate that gastrin deficiency expands CCK2R⁺ stem cells through altered division patterns while priming them for inflammatory plasticity, and that carcinogenic stress drives their transition toward a dysplasia-prone progenitor state.

### A sensory–cholinergic ACh–NGF circuit links G-cell loss to CCK2R⁺ stem-cell expansion

Recent studies by our group have indicated that gastric regeneration and tumorigenesis is also regulated by the peripheral nervous system and specifically the vagus nerve.^22^ Our previous work demonstrated that vagal sensory afferents may sense corpus mucosal injury and trigger a reflex cholinergic response via M3-receptor (M3R) activation, forming a feedback circuit essential for corpus progenitor regeneration.^20^ We hypothesized that a similar neural mechanism operates in the antrum, where gastrin deficiency may activate CCK2R⁺ sensory neurons in the nodose ganglion (NG) to regulate epithelial stem cells.

To trace the antral sensory pathway, we injected the retrograde tracer Fast Blue into the antrum. Five days later, labeled neurons were detected in the NG, confirming that antral afferents project to the NG (Figure 2A). Under homeostasis, a subset of NG neurons expressed CCK2R, partially TRPV1⁺ and CGRP⁺ (Figure 2B and S2A-C). G-cell ablation markedly increased the number of TRPV1⁺CCK2R⁺ and CGRP⁺CCK2R⁺ neurons and induced pERK activation in the NG, while gastrin infusion reversed these effects (Figure 2B, 2C and S2A-D). In parallel, c-FOS expression was elevated in both the nucleus tractus solitarius (NTS) and the dorsal motor nucleus of the vagus (DMV) after G cell ablation, indicating that loss of gastrin activates a sensory–motor vagal reflex (Figure 2D).

**Figure 2.**
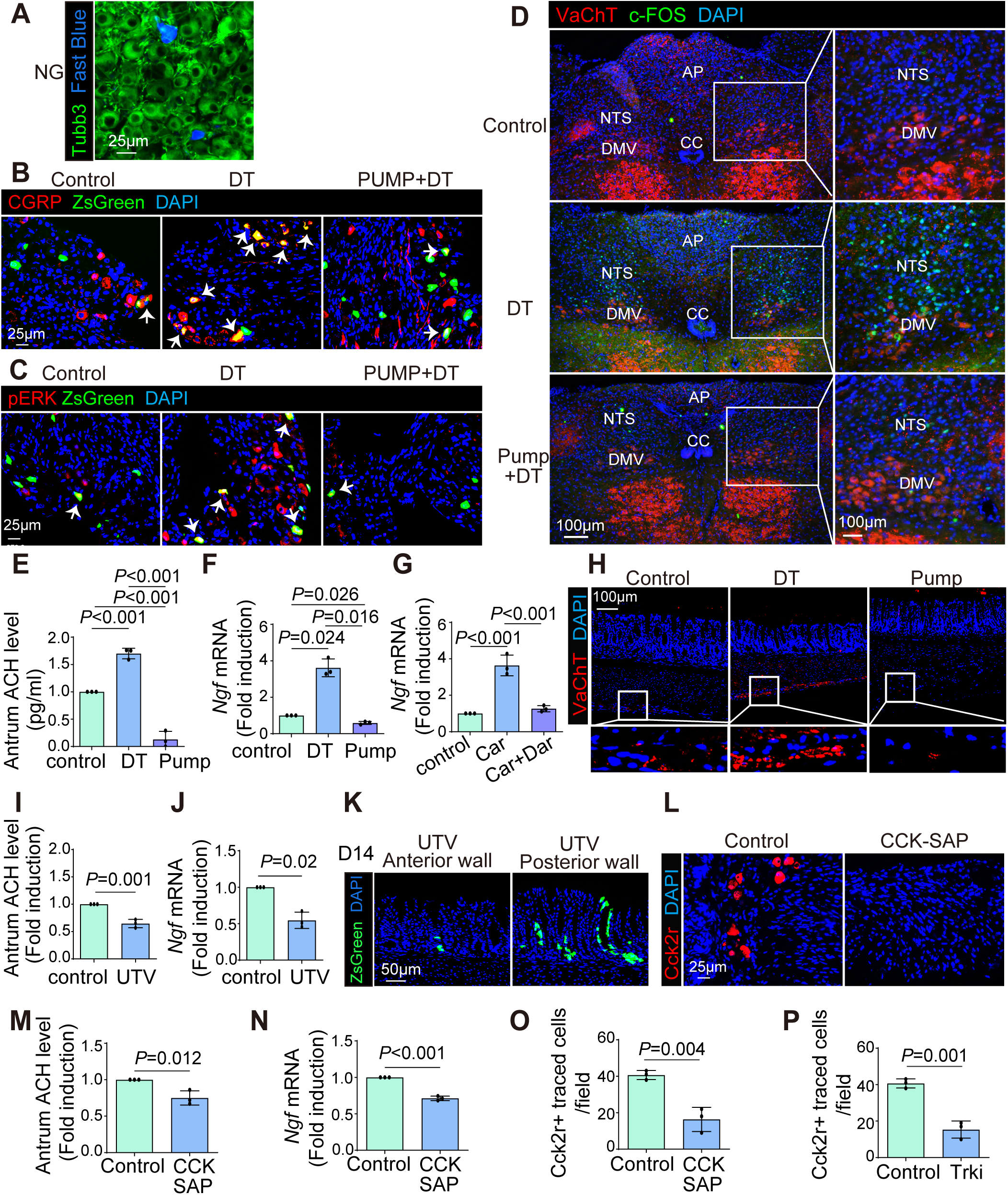
G-cell loss activates a sensory–vagal cholinergic circuit that enhances antral NGF signaling and Cck2r⁺ stem-cell expansion. (A) Retrograde tracing from the antrum identifying Fast Blue⁺ sensory neurons co-labeled with TUBB3 in the nodose ganglion (NG). (B, C) Co-immunofluorescence of ZsGreen with CGRP (B) or pERK (C) in the NG. (D) Co-immunofluorescence of VAChT and c-FOS in the medulla oblongata showing neuronal activation in the nucleus tractus solitarius (NTS) and dorsal motor nucleus of the vagus (DMV). (E) Antral tissue acetylcholine (ACh) levels measured by ELISA. (F) qRT–PCR quantification of *Ngf* mRNA in FACS-sorted antral fibroblasts 2 weeks after treatment. (G) qRT–PCR quantification of Ngf mRNA in cultured FACS-sorted antral fibroblasts under the indicated media conditions. (H) Immunofluorescence images of VAChT staining in the antrum at 9 months. (I-K) Antral ACh levels (I), *Ngf* mRNA levels (J) and Cck2r⁺ lineage tracing (K) 14 days after unilateral truncal vagotomy (UTV). (L–O) Validation of Cck2r⁺ neurons 7 days after CCK-SAP injection (L) and effects on antral ACh (M), *Ngf* expression (N), and Cck2r⁺ cell expansion (O) at 14 days post-injection. (P) Quantification of Cck2r⁺ traced cells per field 14 days after Trk inhibitor treatment. All experiments were performed in Cck2r-CreERT2; Rosa26-ZsGreen; Gastrin-DTR-p2A-TdTomato mice. Data are mean ± SD (n ≥ 3). Scale bars: 25 µm (A, B, C, L), 50 µm (K) and 100 µm (D, H). ACh, acetylcholine; NG, nodose ganglion; UTV, unilateral truncal vagotomy.

Functionally, this circuit activation increased acetylcholine (ACh) release in the antrum (Figure 2E) and was accompanied by a pronounced rise in nerve growth factor (NGF) in antral PDGFRA⁺ fibroblasts isolated by FACS, which remained elevated for up to 9 months (Figure 2F and S2E). Consistent with a direct cholinergic input into this stromal compartment, qRT-PCR showed that Chrm3 (M3R) is the predominant muscarinic receptor expressed in antral PDGFRA⁺ fibroblasts (Figure S2F). In vitro, the cholinergic agonist carbachol robustly induced *Ngf* transcripts in cultured, FACS-sorted PDGFRA⁺ fibroblasts, and this induction was abrogated by the M3R-selective antagonist darifenacin after 48 h of treatment (Figure 2G). Finally, immunostaining for vesicular acetylcholine transporter (VAChT) revealed a marked increase in the density of VAChT⁺ cholinergic fibers following prolonged G-cell ablation, supporting a feed-forward model in which M3R-dependent NGF production by fibroblasts promotes sustained cholinergic remodeling and reinforced antral innervation (Figure 2H and S2G).

To functionally interrogate this neuro–endocrine circuit, we first performed unilateral truncal vagotomy (UTV), which selectively denervates one side of the stomach while preserving overall physiological function.^18,23^ UTV markedly reduced vagal fiber density in the antrum, diminished ACh and NGF levels, and significantly attenuated CCK2R⁺ lineage tracing in the ipsilateral anterior antral wall (Figure 2I-K and S2H, S2I), indicating that intact vagal input is required for CCK2R⁺ stem-cell activation. To establish a direct role for CCK2R⁺ sensory afferents, we next locally injected CCK–saporin (CCK-SAP)^24^ into the right NG to selectively ablate CCK2R⁺ sensory neurons. This manipulation led to a pronounced reduction in antral ACh and NGF levels, accompanied by a significant decrease in Cck2r⁺ lineage expansion (Figure 2L-O and S2J), demonstrating that CCK2R⁺ vagal afferents are necessary to sustain cholinergic output and stem-cell activation. Consistent with a role for NGF-dependent neural reinforcement, pharmacologic inhibition of Trk signaling with selitrectinib for two weeks partially suppressed Cck2r⁺ lineage tracing (Figure 2P and S2K). Together, these complementary loss-of-function approaches establish that the NG–vagal–NGF axis is functionally required for cholinergic-dependent activation and expansion of antral CCK2R⁺ stem cells. Overall, these data establish a gastrin-regulated sensory–cholinergic feedback circuit—linking G-cell loss to NG activation, DMV stimulation, cholinergic outflow, and NGF-mediated reinforcement of Cck2r⁺ stem-cell expansion.

### Loss of G cells promotes expansion of antral Cck2r⁺ stem cells through M3R signaling

To test whether the sensory–cholinergic axis engages antral stem cells through muscarinic signaling, we interrogated our scRNA-seq dataset for muscarinic receptor expression within CCK2R⁺ cells. Chrm3 was selectively enriched, whereas Chrm1 was expressed at low levels and other muscarinic receptor genes were essentially undetectable (Figure S3A and S3B). Notably, Chrm3 expression was further concentrated in the stress-activated, progenitor-like cluster (Figure S3C and S3D), pinpointing a gastrin-sensitive, cholinergic-responsive subpopulation primed for injury-associated reprogramming. To validate these findings, we isolated CCK2R⁺ stem cells from Cck2r-CreERT2; Rosa26-ZsGreen mice by FACS one day after tamoxifen induction and quantified muscarinic receptor transcripts by qRT-PCR. Among all receptor subtypes, Chrm3 was the predominant transcript in antral CCK2R⁺ stem cells (Figure 3A).

**Figure 3.**
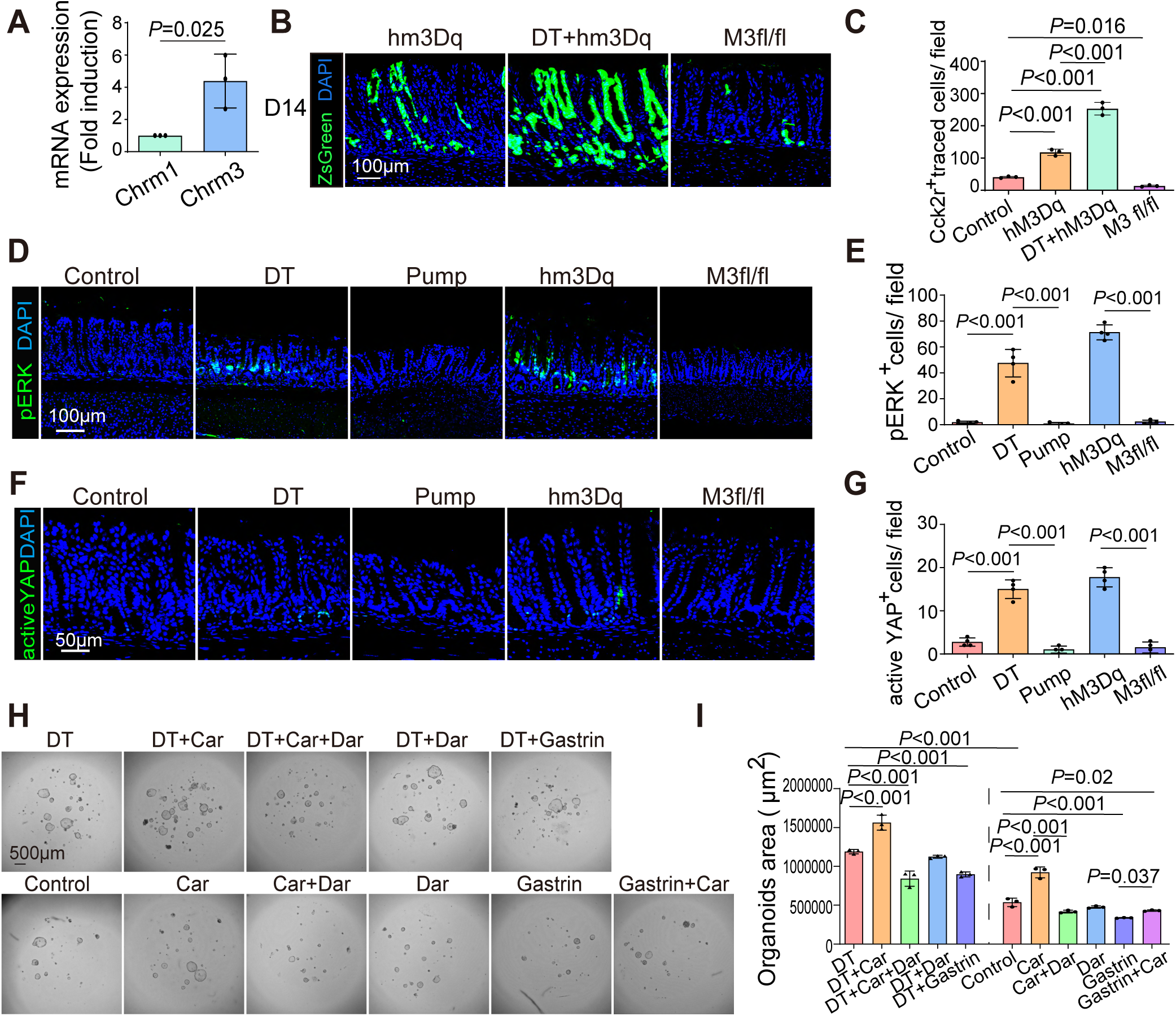
G-cell loss enhances M3R-mediated ERK and YAP signaling to promote Cck2r⁺ stem-cell proliferation. (A) qRT–PCR analysis of muscarinic receptor transcripts (*Chrm1* and *Chrm3*) in FACS-sorted antral Cck2r⁺ cells 1 day after tamoxifen induction. (B, C) Representative images (B) and quantification (C) of Cck2r⁺ lineage tracing 14 days after tamoxifen. (D, E) pERK immunofluorescence (D) and quantification (E) in the antrum at 9 months. (F, G) Active YAP immunofluorescence (F) and quantification (G) in the antrum at 9 months. (H, I) Representative organoids (H) and quantification of organoid formation (I) from sorted single Cck2r⁺ cells after 6 days in culture under the indicated treatments. Data are mean ± SD (n ≥ 3). Scale bars: 100 µm (B, D), 50 µm (F), and 500 µm (H). All experiments were performed in Cck2r-CreERT2; Rosa26-ZsGreen; Gastrin-DTR-p2A-TdTomato mice. Car, carbachol; Dar, darifenacin; DT, diphtheria toxin; Pump, continuous gastrin infusion.

To test the functional relevance of M3R signaling, we used chemogenetic and genetic models. Chemogenetic activation of the M3R in CCK2R^+^ cells through CNO treatment of Cck2rCreERT2; LSL-hM3Dq mice led to a robust increase in lineage tracing, further enhanced by G-cell ablation (Figure 3B and 3C). Of note, this increase was likely due in part to M3R activation in CCK2R^+^ antral stem cells but may also have been driven by the activation of CCK2R^+^ sensory neurons, leading to increased vagal cholinergic tone. However, M3R deletion in Cck2r-CreERT2; Chrm3^fl/fl^ mice strongly reduced lineage tracing, demonstrating that M3R activation is necessary for Cck2r⁺ stem-cell expansion in vivo.

M3R activation is known to engage multiple signaling cascades, including MAPK/ERK,^25^ Akt,^26^ RhoA,^27^ and YAP pathways.^15^ Consistent with these reports, G-cell ablation increased phosphorylated ERK (pERK) and active YAP in the antrum, effects that were further amplified by hM3Dq activation but reversed by continuous gastrin infusion. Conversely, Chrm3 deletion abolished both pERK and YAP activation (Figure 3D-G).

To directly evaluate the downstream effects of M3R signaling on stem-cell behavior, we performed 3D organoid assays using sorted antral CCK2R⁺ cells. Carbachol, a cholinergic agonist, significantly increased organoid size (*P* < 0.05), an effect potentiated by G-cell ablation and blocked by the M3R-selective antagonist darifenacin. In contrast, gastrin supplementation suppressed organoid growth (Figure 3H–I). Importantly, carbachol failed to fully rescue the gastrin-mediated suppression, suggesting that gastrin restrains CCK2R⁺ stem-cell proliferation through both direct epithelial and indirect neural mechanisms.

Mechanistically, carbachol-induced organoid growth was partially reversed by the MEK inhibitor trametinib and the YAP–TEAD inhibitor verteporfin, indicating that M3R signaling promotes stem-cell proliferation through parallel activation of the ERK and YAP pathways (Figure S3E). These results are consistent with the in vivo increase of pERK and nuclear YAP observed in G-cell–ablated mice. Together, these results establish that G-cell loss enhances vagal cholinergic signaling, activating M3R-driven ERK/YAP signaling in antral CCK2R⁺ stem cells to promote their proliferative expansion. Conversely, gastrin restrains this neural– epithelial signaling loop by suppressing both Chrm3 expression and cholinergic input.

### G-cell loss amplifies inflammation and dysplasia in response to chronic antral injury

To determine how gastrin shapes antral epithelial and stem-cell responses to infectious injury, we established long-term *H. pylori* infection in Cck2r-CreERT2; Rosa26-ZsGreen; Gastrin-DTR mice (Figure 4A). Infection was confirmed at 3 and 9 months (Figure 4B and S4A). In control animals, *H. pylori* induced progressive antral atrophy,^28^ beginning at 3 months and worsening by 9 months (Figures 4C, 4D, and S4B-D), accompanied by an initial rise in circulating gastrin followed by marked G-cell depletion and hypogastrinemia (Figure 4E and S4E–H). G-cell ablation further increased bacterial colonization and significantly worsened atrophy scores, whereas both early and delayed gastrin infusion preserved glandular architecture. Across all groups, infection induced inflammatory activation–including higher histologic inflammation scores, increased CD45⁺ infiltration, and upregulation of *Il1b*, *Il6*, *Tnfα* and *Cd45* (Figure 4F–H and S4I–T). These responses were markedly amplified by G-cell loss but attenuated by gastrin supplementation, demonstrating that gastrin constrains mucosal immune tone during chronic infection.

**Figure 4.**
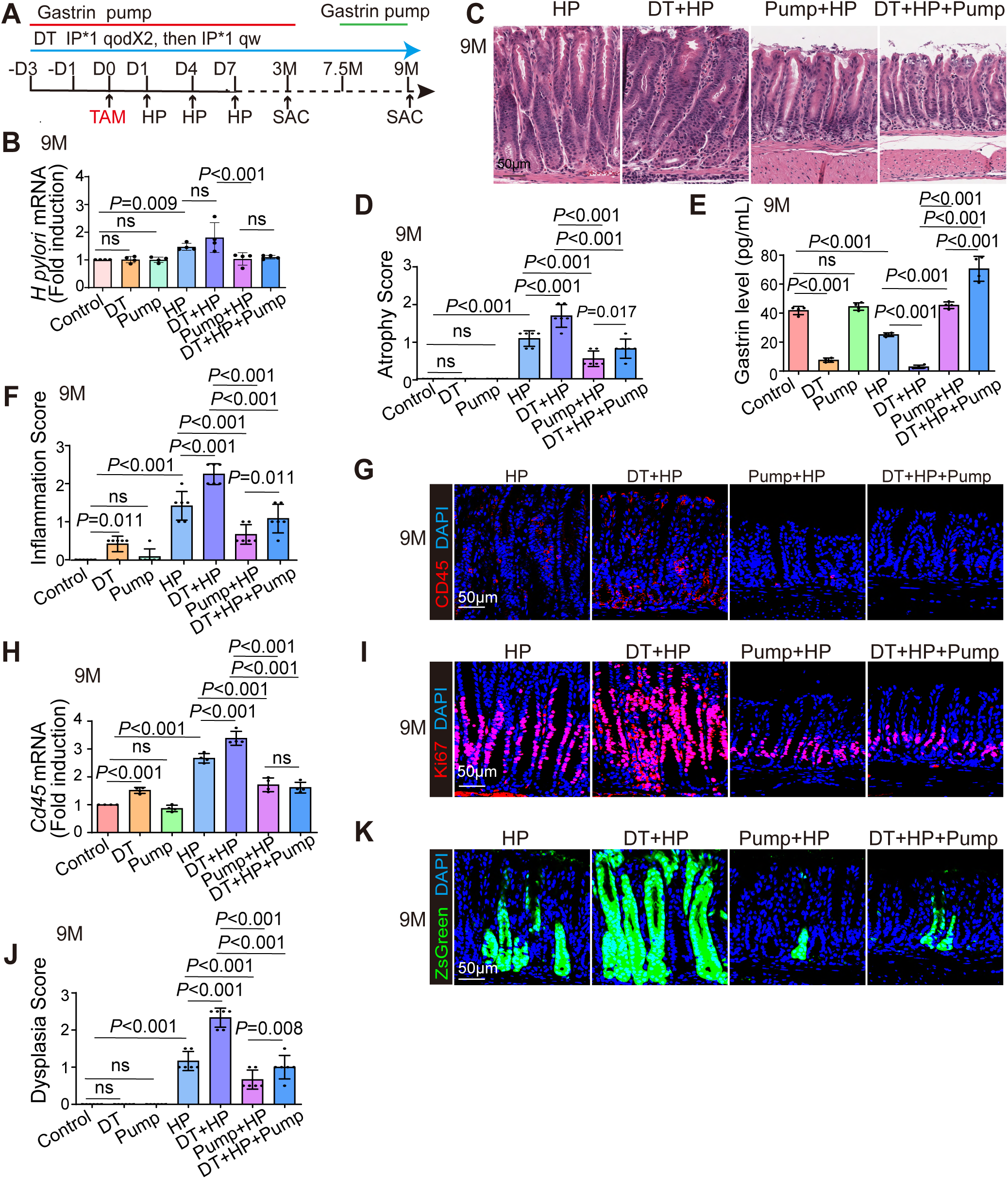
G-cell loss amplifies *H. pylori*-induced antral pathology through enhanced stem cell activation. (A) Experimental scheme of chronic *H. pylori* (Hp) infection. (B–F) *H. pylori* mRNA expression(B), H&E staining (C), atrophy score (D), plasma gastrin (E), and inflammation score (F) of mouse antrum 9 months post-infection. (G and H) CD45⁺ immune cell infiltration via immunofluorescence (G) and mRNA (H) at 9 months. (I and J) Ki67⁺ proliferation (I) and histological dysplasia scores (J) at 9 months. (K) Cck2r⁺ lineage tracing 9 months after tamoxifen induction. Scale bars: 50 µm. Unless otherwise indicated, mice are Cck2r-CreERT2; Rosa26-ZsGreen; Gastrin-DTR-p2A-TdTomato. Hp, *Helicobacter pylori*; DT, diphtheria toxin; Pump, gastrin infusion.

*H. pylori* also increased epithelial proliferation (Ki67⁺ cells and *Ki67* mRNA) (Figure 4I and S5A–F), with G-cell ablation exaggerating and gastrin infusion suppressing this response. By 9 months, infected mice developed mild epithelial dysplasia characterized by glandular distortion, nuclear enlargement, and focal Trop2⁺ staining–an established marker of dysplastic transformation^29^ (Figure 4J and S5G–K). Dysplasia became more extensive and atypical in G-cell–ablated animals but was substantially reduced by gastrin supplementation. Functionally, *H. pylori* expanded CCK2R⁺ lineage–traced clones at both 3 and 9 months (Figure 4K and S5L–N), an effect enhanced by hypogastrinemia and suppressed by hypergastrinemia.

To examine how gastrin signaling influences responses to genotoxic injury, we next employed the N-nitroso-N-methylurea (MNU) model in the same mice (Figure 5A). MNU was given in the drinking water every other week over 10 weeks and at 18 weeks, mice exhibited mild antral inflammation and focal dysplasia, which were strongly exacerbated by G-cell ablation but attenuated by gastrin infusion (Figure S6A–C). Notably, G-cell–ablated mice developed inflammatory infiltration and epithelial abnormalities even without MNU, indicating that hypogastrinemia alone primes the antrum for pathological remodeling. By 9 months, MNU– treated animals exhibited reduced G-cell numbers and hypogastrinemia (Figure 5B and S6D– F), along with moderate inflammation–CD45⁺ immune-cell accumulation and induction of *Cd45*, *Il1b*, *Il6* and *Tnfα*—and elevated epithelial proliferation (Figure 5C–F and S6G–M). These injury responses were again intensified by G-cell loss but were markedly reversed by hypergastrinemia.

**Figure 5.**
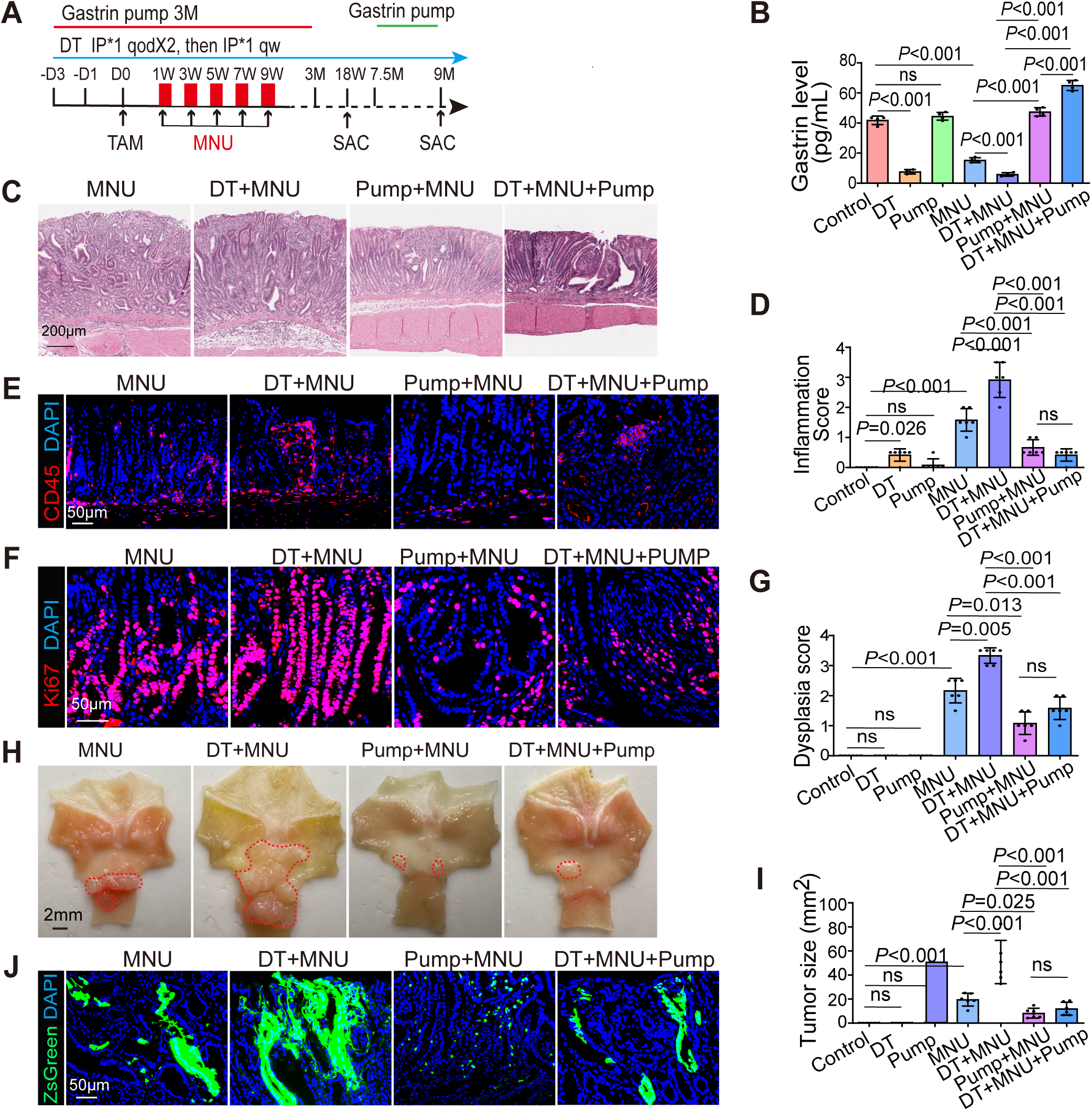
Chronic G-cell loss sensitizes the antrum to inflammation and tumorigenesis post-MNU injury. (A) MNU model scheme. (B) Plasma gastrin levels 9 months post-treatment. (C and D) H&E images (C) and inflammation scores (D) after 9 months. (E–G) CD45⁺ infiltration (E), Ki67⁺ proliferation (F), and dysplasia scores (G) at 9 months. (H and I) Gross images (H) and tumor burden quantification (I) 9 months post-MNU. (J) Cck2r⁺ lineage tracing 9 months after tamoxifen induction. Scale bars: 2 mm (H), 200 µm (C), 50 µm (E, F, J). Unless otherwise indicated, mice are Cck2r-CreERT2; Rosa26-ZsGreen; Gastrin-DTR-p2A-TdTomato. MNU, N-nitroso-N-methylurea. DT, diphtheria toxin; Pump, gastrin infusion.

When carried out to 9 months, mice treated with MNU exhibited grossly visible antral tumors (Figure 5C, G–I and S6N). Tumor burden and dysplasia severity were substantially increased in G-cell–ablated mice, whereas both early and delayed gastrin infusion reduced tumor size and histologic atypia. Lineage tracing revealed robust tracing of tumors by CCK2R⁺ stem-cell– derived clones after MNU injury, which was further enhanced by hypogastrinemia but suppressed by hypergastrinemia (Figure 5J and S6O). Together, these results show that chronic gastrin deficiency sustains inflammatory signaling and drives hyperactivation of the CCK2R⁺ stem-cell compartment, thereby heightening susceptibility to dysplastic transformation of CCK2R+ progenitors under chronic infectious or genotoxic stress.

### G-cell loss promotes M3R-mediated cholinergic activation and parallel ERK and YAP signaling in antral tumorigenesis

Given that NGF expression and vagal cholinergic activity was elevated following G-cell loss, we next examined whether this neural remodeling persists under carcinogenic stress. MNU exposure alone increased *Ngf* mRNA expression and VAChT⁺ cholinergic fiber density in the antrum, and this effect was further amplified by G-cell ablation (DT+MNU). In contrast, continuous gastrin infusion markedly reduced *Ngf* mRNA level and cholinergic innervation in MNU-treated mice (Figure 6A, 6B and S7A). qRT-PCR analysis confirmed robust *Chrm3* induction following prolonged G-cell loss and carcinogenic stress, with the highest expression in DT+MNU mice, thus correlating with hypogastrinemia plus genotoxic stress, whereas gastrin supplementation downregulated *Chrm3* expression even at late stages (Figure 6C). Together, these data indicate that gastrin deficiency enhances vagal cholinergic signaling and epithelial M3R expression under carcinogenic stress, while gastrin counteracts this activation by dampening both vagal output and receptor abundance.

**Figure 6.**
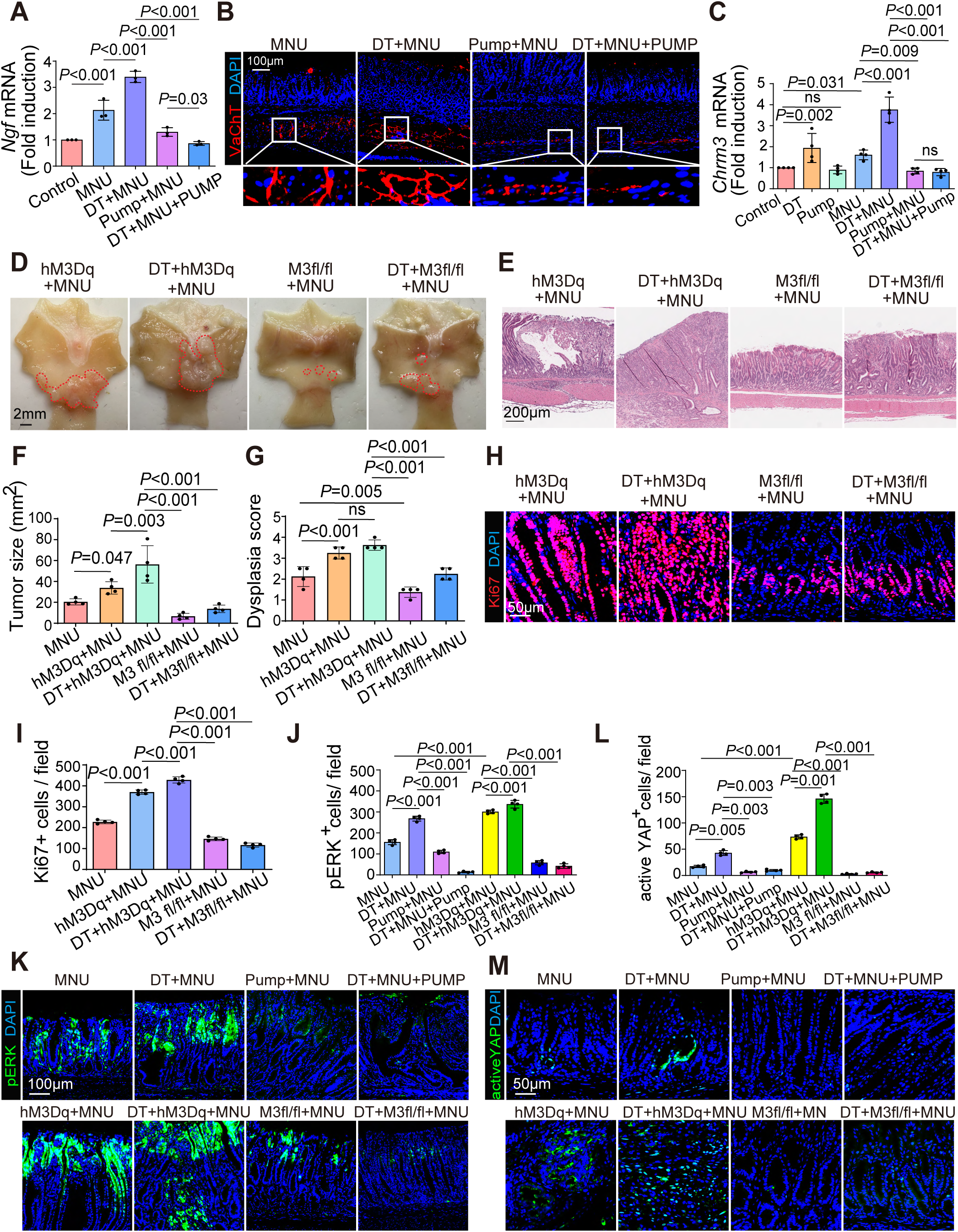
G-cell loss enhances vagal cholinergic remodeling and M3R signaling to drive antral inflammation and dysplasia. (A) qPCR analysis of *Ngf* expression in antral tissue after 9 months. (B) Representative immunofluorescence of VAChT⁺ cholinergic fibers in the antrum after 9 months. (C) qPCR analysis of *Chrm3* expression in antral tissue after 9 months. (D) Representative gross images of MNU-induced antral tumors. (E and F) Representative H&E staining (E) and quantification of antral tumor size (F) after 9 months of MNU treatment. (G) Histological dysplasia scores in the antrum after 9 months. (H and I) Representative immunofluorescence image (H) and quantification (I) of Ki67⁺ epithelial proliferation. (J and K) Quantification (J) and representative image (K) of phosphorylated ERK (pERK) in the antrum. (L and M) Quantification (L) and representative image (M) of active YAP expression in the antrum. Data are mean ± SD (n ≥ 3). Scale bars: 2 mm (D), 200 µm (E), 100 µm (B, K), 50 µm (H, M). DT, diphtheria toxin; Pump, continuous gastrin infusion; MNU, N-nitroso-N-methylurea; CNO, clozapine-N-oxide.

Functionally, sustained M3R activation drove tumor progression. In Cck2r-CreERT2; LSL-hM3Dq mice, chronic CNO administration enlarged antral tumors and increased Ki67⁺ proliferation after 9 months of MNU exposure, effects further intensified by G-cell ablation. Conversely, genetic deletion of Chrm3 in Cck2r-CreERT2; Chrm3^fl/fl^ mice markedly reduced tumor burden and proliferation, even under gastrin-deficient conditions (Figure 6D-I), establishing M3R signaling as a key effector of tumorigenesis in hypogastrinemia.

At the signaling level, MNU exposure maintained and further amplified ERK and YAP activation downstream of M3R signaling, consistent with a sustained pro-proliferative state in G-cell–deficient antrum. Chemogenetic activation of M3R (hM3Dq) further enhanced, whereas gastrin infusion or Chrm3 deletion markedly suppressed, both pathways (Figure 6J– M). These findings identify ERK and YAP as parallel downstream effectors of M3R signaling that cooperatively drive the proliferative and tumor-promoting phenotype of CCK2R⁺ stem cells.

To pharmacologically validate the role of M3R signaling in tumor progression, we administered the selective M3R antagonist darifenacin via osmotic pump during the final month of MNU group. Darifenacin treatment significantly reduced overall tumor burden, Ki67⁺ epithelial proliferation and pERK and pYAP expression (Figure S7B-I), demonstrating that late-stage inhibition of M3R signaling reverses tumor-associated hyperproliferation and supports the therapeutic potential of targeting M3R in gastrin-deficient cancer.

In parallel, targeting the neural NGF–TrkA axis produced similar effects. Oral selitrectinib given during the final month of MNU exposure reduced VAChT⁺ fiber density, decreased tumor size, and pERK and pYAP expression (Figure S7J-Q). Together, these findings demonstrate that both the neural (NGF–TrkA) and epithelial (M3R–ERK/YAP) branches of the vagal cholinergic circuit cooperate to sustain antral tumorigenesis in gastrin-deficient states and are each pharmacologically targetable.

Collectively, these results define a dual neuroepithelial signaling axis–comprising the NGF– TrkA–vagal sensory branch and the M3R–ERK/YAP epithelial branch–that drives tumorigenesis from CCK2R^+^ antral stem cells in the gastrin-deficient antrum and provides multiple points for potential therapeutic intervention.

### Clinical evidence links G-cell loss to CCK2R⁺ stem-cell reprogramming during human antral carcinogenesis

To assess the clinical relevance of our findings, we re-analyzed publicly available single-cell RNA-sequencing data from gastric antral biopsies of nine patients spanning the pathological continuum from chronic gastritis to early gastric cancer (EGC).^30^ Samples were categorized into three stages—gastritis, intestinal metaplasia (IM), and gastric cancer (GC)—to capture progressive epithelial transformation.

Unsupervised clustering identified eight transcriptionally distinct epithelial populations (Figure 7A). G cells declined progressively across disease stages, with a pronounced reduction in IM and near-complete loss of G cells in GC, closely recapitulating the loss of gastrin-expressing cells observed in our mouse model (Figure 7B–C). Notably, a GC-specific epithelial cluster emerged exclusively in tumor samples, displaying the highest CytoTRACE score—comparable to the stress-activated progenitor-like population identified in MNU-treated, G-cell–ablated mice (Figure 1K, 7D)—suggesting conserved transcriptional plasticity between murine and human antral transformation.

**Figure 7.**
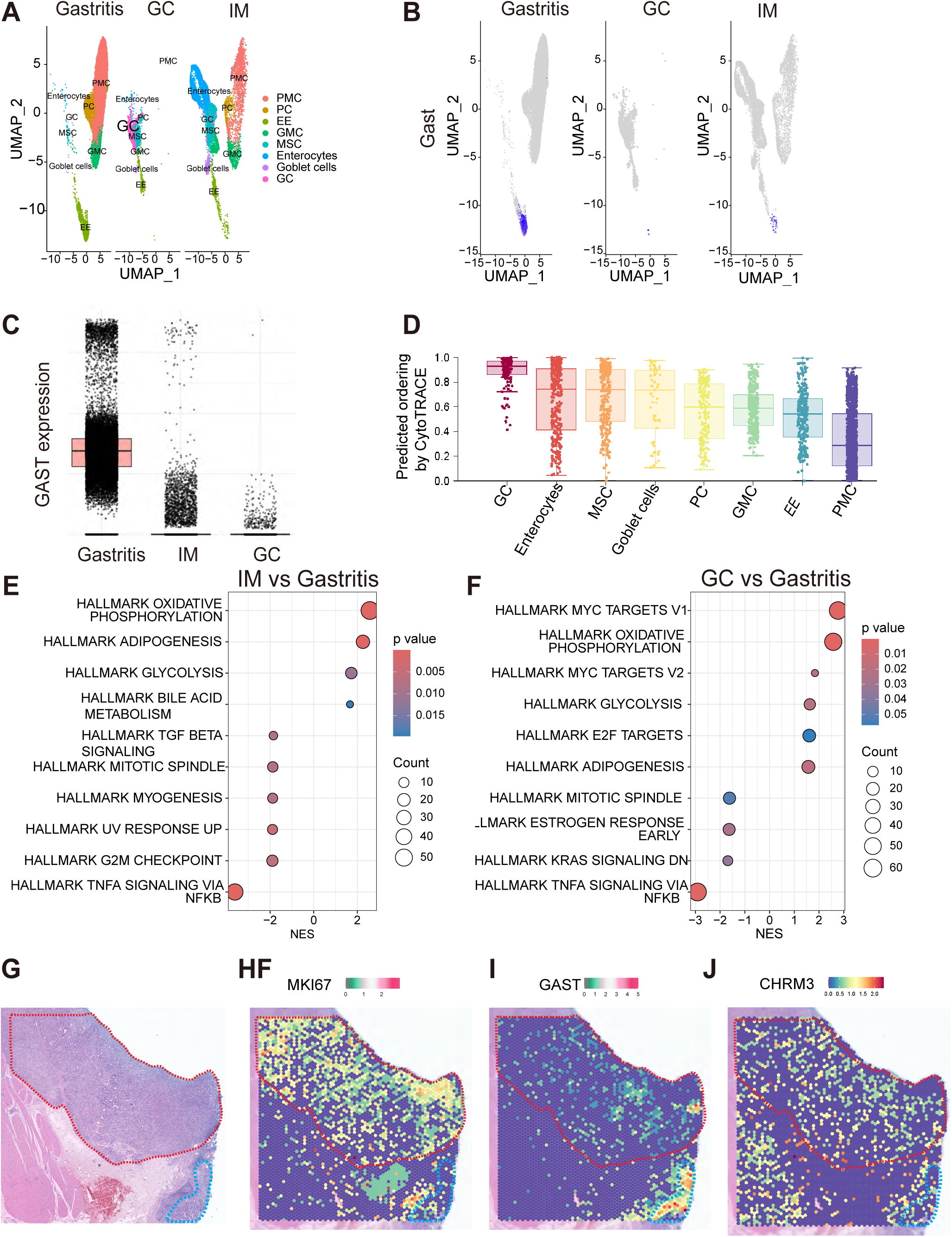
Progressive G-cell loss and CHRM3⁺ progenitor reprogramming during human antral carcinogenesis. (A) UMAP visualization of epithelial cells from human gastric antral biopsies across gastritis, intestinal metaplasia (IM), and gastric cancer (GC) stages. (B and C) UMAP feature plots (B) and quantification (C) demonstrating progressive depletion of *GAST* (gastrin) expression from gastritis to IM and GC. (D) CytoTRACE analysis identifying a GC-associated epithelial cluster with a high progenitor-like transcriptional state. (E and F) Gene set enrichment analysis (GSEA) of hallmark pathways comparing IM (E) or GC (F) versus gastritis. (G) Representative H&E image of human gastric cancer highlighting tumor (red outline) and adjacent normal (blue outline) regions. (H–J) Spatial transcriptomic maps showing increased *MKI67* expression (H) and depletion of *GAST* (I) and increased *CHRM3* (J) within tumor regions relative to adjacent mucosa. IM, intestinal metaplasia; GC, gastric cancer cells; GMC, antral basal gland mucous cells; MSC, metaplastic stem-like cells; PC, proliferative cells; PMC, pit mucous cells; EE, enteroendocrine cells.

Gene set enrichment analysis (GSEA) revealed a metabolic transition during IM, characterized by upregulated oxidative phosphorylation, followed by activation of dysplasia-associated pathways in GC, including MYC signaling and inflammatory reprogramming (Figure 7E–F). These signatures closely paralleled the molecular programs identified in murine CCK2R⁺ progenitors under chronic hypogastrinemia and M3R activation.

Spatial transcriptomic profiling of human gastric antral tumor further demonstrated progressive loss of G cells, accompanied by increased Ki67 and CHRM3 expression within tumor regions compared with adjacent non-neoplastic mucosa (Figure 7G-J). This pattern mirrored the expansion of proliferative, M3R⁺ CCK2R⁺ stem cells in G-cell–ablated mice, reinforcing a conserved gastrin–cholinergic–M3R axis in human gastric carcinogenesis.

Together, these analyses reveal that G-cell depletion, expansion of GC-specific epithelial populations, and transcriptional reprogramming toward a proliferative, inflammatory state are conserved features of human and murine antral tumorigenesis—underscoring gastrin-dependent neuroepithelial regulation as a key barrier against gastric cancer progression.

## DISCUSSION

Our findings uncover a neuro–endocrine surveillance mechanism in the gastric antrum, where the peripheral nervous system detects the loss of the endocrine signal gastrin and converts this information into a pro-regenerative, and ultimately preneoplastic, epithelial program. We show that gastrin deficiency is not a passive hormonal deficit, but a potent activating cue sensed by CCK2R⁺ vagal afferents, which triggers a nodose–NTS–DMV sensory–motor reflex that elevates acetylcholine release, induces sustained NGF–TrkA–dependent cholinergic remodeling, and upregulates epithelial M3R. This neural remodeling drives ERK- and YAP-dependent activation of CCK2R⁺ +4 stem cells and, together with gastrin’s direct suppression of symmetric division, promotes their expansion and the emergence of an injury-responsive progenitor state highly susceptible to inflammation and dysplasia. These data reveal a previously unappreciated endocrine gatekeeping function of G cells, positioning gastrin as a tonic suppressor of vagal circuits that regulate antral stem-cell plasticity.

Although gastrin has long been understood primarily as a regulator of parietal cell function and gastric acid secretion^31^, its trophic actions in the corpus—particularly in promoting CCK2R⁺ progenitors expansion during ulcer repair^20^—suggest broader roles in epithelial homeostasis. In the antrum, early studies using germline Gastrin–/– mice demonstrated that gastrin limits symmetric division of CCK2R⁺ stem cells and restrains tumorigenesis^9^, establishing a foundational link between gastrin and antral stem-cell regulation. However, these models were based on whole-body, lifelong gastrin deficiency and thus conflated developmental effects with adult physiology while implicitly assuming that gastrin acts solely through direct epithelial mechanisms. Such approaches could not determine whether gastrin suppresses neural activity or whether loss of endocrine input activates extrinsic circuits that modulate stem-cell behavior. By employing temporally controlled, adult-onset G-cell ablation, our study demonstrates that gastrin loss is sufficient to activate a CCK2R⁺ vagal reflex and that endocrine deficiency cooperates with neural remodeling to expand the CCK2R⁺ stem-cell pool—clarifying how gastrin status shapes antral regenerative and preneoplastic trajectories.

Our work builds upon and extends the emerging principle that neural circuits regulate epithelial stem cells across diverse tissues. Sensory and sympathetic neurons modulate hematopoietic stem-cell quiescence, melanocyte stem-cell aging, and intestinal stem-cell (ISC) maintenance, while stress-activated central pathways suppress ISC stemness through descending vagal signals.^16,17,32,33^ In the stomach, vagal cholinergic signaling and tuft-cell–derived acetylcholine have been shown to promote epithelial proliferation and tumor growth through muscarinic pathways,^15^ and cancer–nerve interactions contribute to gastric carcinogenesis.^22^ Yet these studies did not define which epithelial populations receive muscarinic input, how neural activity is modulated by endocrine cues such as gastrin, whether vagal sensory afferents respond to endocrine-cell loss, or how neural and endocrine pathways converge at the stem-cell level. Previous work on vagal sensory signaling has focused predominantly on CCKAR-expressing afferents, which robustly sense luminal and enteroendocrine CCK to regulate satiety and brainstem activation.^34,35^ TRPV1 has been shown to potentiate CCK signaling in these neurons, enhancing transmission to the NTS.^36^ By contrast, far less is known about CCKBR (CCK2R)–expressing vagal afferents, which bind gastrin but not CCK, and whose physiological roles have remained largely unexplored. Our findings address this gap by demonstrating that gastrin deficiency is actively sensed by CCK2R⁺ vagal afferents, triggering a nodose–NTS–DMV reflex that amplifies cholinergic output and remodels the antral niche. This reveals a previously unrecognized form of endocrine surveillance, distinct from classical CCKAR-mediated nutrient sensing, through which loss of gastrin engages neural circuits to drive stem-cell reprogramming.

Our data reveal that gastrin loss paradoxically enhances vagal cholinergic sprouting, creating a feed-forward loop in which NGF–TrkA–dependent remodeling further augments vagal output. Prior studies have shown that NGF can be induced by cholinergic stimuli,^37,38^ and in turn promote cholinergic nerve growth,^15,39^ consistent with this circuit-level reinforcement. Within this framework, CCK2R⁺ stem cells emerge as direct epithelial targets of vagal cholinergic signaling through M3R, which we identify as selectively enriched in an injury-responsive progenitor-like subset and strongly induced by gastrin deficiency and carcinogenic stress. M3R has been reported to activate MAPK,^25^ Akt,^26^ YAP,^15^ and RhoA^27^ pathways in various contexts, and our findings position M3R as the principal epithelial effector through which vagal inputs act to activate ERK and YAP in CCK2R⁺ stem cells. Thus, muscarinic signaling dynamically links endocrine deficiency and neural remodeling to stem-cell reprogramming.

The translational relevance of this axis is supported by human scRNA-seq and spatial datasets demonstrating progressive G-cell depletion, and induction of proliferative and inflammatory programs across the gastritis–metaplasia–early cancer continuum. These conserved features suggest that the gastrin–sensory–vagus–M3R axis operates in human gastric disease and that dysregulated neuro–endocrine communication may mark early antral transformation. Although additional cellular contributors—including stromal fibroblasts, immune infiltrates, and enteric glia—are likely to participate in this circuitry, our findings raise the possibility that M3R antagonists, NGF–TrkA inhibitors,^40,41^ or strategies that sustain gastrin tone could modulate this axis to restrain preneoplastic progression in high-risk settings such as chronic *H. pylori* infection or severe antral atrophy with hypogastrinemia.

In summary, we show that the loss of an endocrine cell population is actively sensed by the vagal nervous system and transformed into a stem-cell–activating signal, revealing a previously unrecognized endocrine–neural–epithelial axis that governs antral stem-cell plasticity, inflammation, and preneoplastic evolution. These findings reshape the conceptual framework for how the gastric antral niche integrates hormonal and neural cues and highlight new opportunities for targeted interception of early gastric disease.

### Limitations of the study

While we delineate both direct epithelial effects of gastrin on CCK2R⁺ stem-cell division behavior and indirect neural remodeling that amplifies M3R–ERK/YAP signaling, the relative magnitude and temporal hierarchy of these two arms across diverse injury and inflammatory contexts remain to be determined. In addition, our human data reveal associations between G-cell depletion, cholinergic/NGF signatures, and progenitor activation, but do not establish causality, which will require prospective sampling and functional perturbation in human-derived systems.

## RESOURCE AVAILABILITY

### Lead contact

Requests for further information and resources should be directed to and will be fulfilled by the lead contact, Timothy C. Wang.

### Materials availability

Requests for mouse lines used in this study should be directed to the lead contact and will be fulfilled upon reasonable request. This study did not generate new unique reagents.

### Data and code availability

The scRNA-seq dataset generated during this study is available in the Gene Expression Omnibus (GEO, GSE317614) and Genome Sequence Archive database, and the accession number is listed in the key resources table. This paper does not report original code. Any additional information required to reanalyze the data reported in this paper is available from the lead contact upon request.

## ACKNOWLEDGMENTS

We are thankful for the help from Dr. James G. Fox (MIT) for providing the *H. pylori SS1* strain. We acknowledge the patients and their families for their participation in this study. This research was funded by NIDDK grant R01DK128195 (TCW), NCI grants R35CA210088 (TCW), and R01CA272901 (TCW, JQ); and a Department of Defense grant W81XWH-21-10901 (TCW). QW is supported by NIH F31DK142531. This study was also supported by National Natural Science Foundation of China General Program (Grant No. 82570660, BZ), National Natural Science Foundation of China Young Scientists Fund (Grant No. 82203308, BZ), Fujian Provincial Natural Science Foundation of China (Grant No. 2026J010043, BZ), Fujian Provincial Natural Science Foundation of China (Grant No. 2025J01116, BZ) and the National Key Clinical Specialty Construction Project of Fujian Province, China (Grant No. 2023-1594, FC). This work was also supported by the NIH/NCI Cancer Center Support Grant (P30CA013696) and used the resources of the Herbert Irving Comprehensive Cancer Center, including the Flow Cytometry Shared Resources, Molecular Pathology/MPSR, Genomics and High Throughput Screening, and the Genetically Modified Mouse Model Shared Resource (GMMMSR); as well as the Columbia University Digestive and Liver Disease Research Center (CU-DLDRC) grant P30DK132710 and its Bio-Imaging, Organoid, Biospecimen and Bioinformatics cores. This work was also supported by the Influx sorter: S10OD020056.

## AUTHOR CONTRIBUTIONS

Conceptualization: BZ, HZ, RT, FC, GL, EM and TCW ; methodology: BZ, HZ, HK, RT, JL, KH, XZ, GL, FW, YO, QW, EM, YZ, PZ, SL, LZ, XW, FC, XX, JA and TCW; software and formal analysis: BZ, HK, JQ, JuL, MH, YZ and TCW; investigation: BZ, HK, RT, KH; writing– original draft: BZ; writing–review and editing: BZ, HZ, TC, FC and TCW; supervision: TCW. All authors read and approved of the final manuscript. TCW is the guarantor.

## STAR★METHODS

Detailed methods are provided in the online version of this paper and include the following:

● KEY RESOURCES TABLE
● EXPERIMENTAL MODEL AND SUBJECT DETAILS

○ Animal models
○ Mouse surgery
● METHOD DETAILS

○ Ethics approval
○ Chronic H. pylori infection model and MNU-induced injury model
○ H&E staining
○ Immunofluorescence
○ Histological evaluation
○ Gastrin Quantification
○ Enzyme-linked immunosorbent assay (ELISA)
○ Antrum dissection and single-cell isolation
○ Flow cytometry and cell sorting
○ Fibroblast cells culture
○ 3D Culture
○ RNA extraction and qRT-PCR
● QUANTIFICATION AND STATISTICAL ANALYSIS

○ Mouse Single Cell RNA-seq Analysis
○ Human Single Cell RNA-seq Analysis
○ Processing of spatial transcriptomics data
○ Statistical analysis

Supplemental Table 1. Top 50 marker genes for each cluster in mouse antral scRNA-seq.

Supplemental Table 2. Sequences of qRT-PCR Primers, related to STAR Methods.

## STAR★METHODS

### KEY RESOURCES TABLE

| REAGENT or RESOURCE | SOURCE | IDENTIFIER |
| --- | --- | --- |
| Antibodies |  |  |
| Mouse anti-Ki67 | BD Pharmingen | Cat# 550609, RRID: AB_393778 |
| Goat anti-TROP-2 | R&D Systems | Cat# AF1122, RRID: AB_2205662 |
| Rabbit anti-Gastrin | Dako | Cat# A0568, RRID: N/A |
| Rabbit anti-CD45 | Abcam | Cat# ab10558, RRID: AB_442810 |
| Rabbit anti-VACHT | Synaptic Systems | Cat# 139103, RRID: AB_887864 |
| Rabbit anti-ZsGreen | Takara Bio | Cat# 632474, RRID: AB_2491179 |
| Goat anti-tdTomato | Biorbyt | Cat# orb182397 , RRID: AB_2687917 |
| Guinea pig anti c-Fos | Synaptic Systems | Cat# 226308, RRID: AB_2905595 |
| TUBB3 | Abcam | Cat# ab78078, RRID: AB_2256751 |
| goat anti-CGRP | Abcam | Cat# ab36001, RRID: AB_725807 |
| Rabbit anti-TRPV1 | Alomone Labs | Cat# ACC-030, RRID: AB_2313819 |
| Rabbit anti-Phospho-Erk1/2 | Cell Signaling Technology | Cat# 4370S, RRID: AB_2315112 |
| Rabbit anti-active YAP1 | Abcam | Cat# ab205270, RRID: AB_2813833 |
| APC anti-human CD140a (PDGFRα) Antibody (Clone: 16A1) | BioLegend | Cat# 323511, RRID: AB_2783190 |
| BD OptiBuild™ BV650 Rat Anti-Mouse CD31 | Fisher Scientific | Cat# 740483, RRID: AB_2740207 |
| Goat anti-Rabbit IgG (H+L) Alexa Fluor 488 | Invitrogen | Cat# A11008, RRID: AB_143165 |
| Goat anti-Rabbit IgG (H+L) Alexa Fluor 555 | Invitrogen | Cat# A21428, RRID: AB_141784 |
| Goat anti-mouse IgG (H+L) Alexa Fluor 555 | Thermo Fisher Scientific | Cat# A21422, RRID: AB_2535844 |
| Donkey anti-goat IgG (H+L) Alexa Fluor 488 | Thermo Fisher Scientific | Cat# A32814, RRID: AB_2762838 |
| Donkey anti-goat IgG (H+L) Alexa Fluor 555 | Thermo Fisher Scientific | Cat# A32816, RRID: AB_2762839 |
| Goat anti-Guinea Pig (Alexa Fluor 488) | Thermo Fisher Scientific | Cat# A-11073, RRID: AB_2534117 |
| Brilliant Violet 785™ anti-mouse CD326 (Ep-CAM) Antibody | BioLegend | Cat# 118245, RRID: AB_2860639 |
| APC anti-mouse CD326 (Ep-CAM) | BioLegend | Cat# 118214, RRID: AB_1134102 |
| Biological samples |  |  |
| *H. pylori* SS1 strain | Dr. James G. Fox (MIT) | NA |
| Chemicals, peptides, and recombinant proteins |  |  |
| Tamoxifen | Sigma-Aldrich | Cat# T5648 |
| Diphtheria toxin (DT) | EMD Millipore | Cat# 322326 |
| DPBS | Fisher Scientific | Cat# 14-190-250 |
| Ultra Pure 0.5M EDTA, pH 8.0 | Invitrogen | Cat# 15575020 |
| TrypLE Express | Thermo Fisher Scientific | Cat# 12-604-021 |
| Dnase I | Thermo Fisher Scientific | Cat# 10104159001 |
| Advanced DMEM/F12 | Gibco | Cat# 12634-010 |
| GlutMAX | Thermo Fisher Scientific | Cat# 35050061 |
| HEPES | Thermo Fisher Scientific | Cat# 15-630-080 |
| Antibiotic-Antimycotic | Thermo Fisher Scientific | Cat# 15240062 |
| Fetal bovine serum (FBS) | Gemini Bio-Products | Cat# 900-108 |
| B27 supplement | Thermo Fisher Scientific | Cat# 17504044 |
| N-2 supplement | Fisher Scientific | Cat# 17502048 |
| N-Acetyl-L-cysteine | Sigma-Aldrich | Cat# A9165 |
| EGF | Thermo Fisher Scientific | Cat# PMG8043 |
| Human Heat Stable bFGF Recombinant Protein | Fisher Scientific, | Cat# PHG0369 |
| Y-27632 (dihydrochloride) | MedChem Express | Cat# HY-10583 |
| Gastrin I (human) | Tocris | Cat# 3006 |
| Darifenacin hydrobromide | Sigma-Aldrich | Cat# SML1102 |
| Fast blue | Polysciences | Cat# 17740-1 |
| CCK-SAP | Advanced Targeting Systems | Cat# KIT-31 |
| Clozapine N-oxide dihydrochloride | Fisher Scientific | Cat# 6329 |
| Selitrectinib (LOXO-195) | Ambeed Inc | Cat# A285925 |
| Carbachol | EMD Millipore | Cat# 212385 |
| Trametinib | Cayman Chemical | Cat# 16292 |
| Verteporfin | MedChem Express | Cat# HY-B0146 |
| DAPI Solution | BD Biosciences | Cat# 564907 |
| 32% Paraformaldehyde Aqueous Solution, EM Grade | Electron Microscopy Sciences | Cat# 15714 |
| Signalstain® Antibody Diluent | Cell Signaling | Cat# 8112L |
| Corning® Matrigel® Growth Factor Reduced (GFR) Basement | Corning | Cat# 356231 |
| VECTASHIELD® Antifade Mounting Medium, With DAPI | Vector Laboratories | Cat# H-1200-10 |
| Brucella broth | Fisher Scientific | Cat# 211088 |
| N-nitroso-N-methylurea | Gojira Fine Chemicals | Cat# NM 1003 |
| Antigen Unmasking Solution, Tris-Based | Vector Laboratories | H-3301 |
| Antigen Unmasking Solution, Citrate-Based | Vector Laboratories | H-3300 |
| Qiagen RNA isolation kit | Qiagen | Cat# 74034 |
| qScript cDNA SuperMix | QuantaBio | Cat# 95048 |
| SYBR Green qPCR mix | Quantabio | Cat# 101414-286 |
| Primers for qRT-PCR, see Table S1 | This paper | NA |
| Critical commercial assays |  |  |
| ACh ELISA Kit | Elabscience | Cat# E-EL-0081 |
| Gastrin EIA Kit | Sigma-Aldrich | Cat# RAB0200 |
| Deposited data |  |  |
| Singe-cell RNA-seq data of CCK2R+ cells from mouse antrum | This paper | GEO: GSE317614 |
| Singe-cell RNA-seq data of human gastric premalignant lesions and early gastric cancer | Zhang et al.<sup>30</sup> | GEO: GSE134520 |
| Spatial transcriptomics data of human gastric cancer | Wang et al.<sup>42</sup> | Genome Sequence Archive database (accession numbers: HRA011663) |
| Experimental models: Cell lines |  |  |
| L-WRN cells | Adam Bass (Novartis) | N/A |
| Experimental models: Organisms/strains |  |  |
| Mouse C57BL/6J | The Jackson Laboratory | Stock# 000664 |
| Mouse C57BL/6J Lgr5-DTR-GFP | Tian et al.<sup>43</sup> | NA |
| Mouse C57BL/6J Gastrin-DTR-p2A-TdTomato | Zheng et al.<sup>20</sup> | NA |
| Mouse C57BL/6J Cck2r-CreERT2-BAC | Hayakawa et al.<sup>8</sup> | NA |
| Mouse Rosa26-tdTomato: Gt(ROSA)26Sortm9(CAG-tdTomato)Hze/J | The Jackson Laboratory | Stock# 007909 |
| Mouse B6.Cg-Gt(ROSA)26Sortm6(CAG-ZsGreen1)Hze/J | The Jackson Laboratory | Stock# 007906 |
| Mouse LSL-hM3Dq | The Jackson Laboratory | Stock# 026220 |

| Software and algorithms |  |  |
| --- | --- | --- |
| ImageJ version1.6/Fiji | NIH | http://imagej.net |
| FlowJo V10 | FlowJo | https://www.flowjo.com |
| 10X CellRanger software: version 3.1 | 10X Genomics | https://support.10xgenomics.com/singlecell-gene-expression |
| R-4.4.1 | Bell Laboratories | https://www.r-project.org/about.html |
| GraphPad prism 9.1.2 | GraphPad Software | http://graphpad.com |
| Seurat R package (version 5.1.0) | Hao *et al.*<sup>44</sup> | https://satijalab.org/seurat/ |
| COSG package (version 0.9.0) | Dai *et al.*<sup>45</sup> | https://github.com/genecell/COSG |
| CellMarker 2.0 | Hu *et al.*<sup>46</sup> | http://117.50.127.228/CellMarker/index.html |
| inferCNV (v.1.20.0) | Patel *et al.*<sup>47</sup> | https://github.com/broadinstitute/inferCNV |
| Monocle3 | Cao *et al.*<sup>48</sup> | https://cole-trapnell-lab.github.io/monocle3/ |
| CytoTRACE | Gulati *et al.*<sup>49</sup> | https://cytotrace.stanford.edu |
| Space Ranger (v1.3.0) | 10X Genomics | https://www.10xgenomics.com/support/software/space-ranger/ |
| stLearn | Pham *et al.*<sup>50</sup> | https://stlearn.readthedocs.io/en/stable/index.html |
| EndNote X9 | Clarivate | https://support.clarivate.com/Endnote/s/article/EndNote-Download-link-for-older-EndNote-versions?language=en_US |
| Other |  |  |
| Alzet osmotic pump | Alzet | Cat# 1007D, 1002,<br>1004, 2006 |

### EXPERIMENTAL MODEL AND SUBJECT DETAILS

#### Animal models

All animal experiments and procedures were conducted following the NIH guidelines for animal research and were reviewed and approved by the Institutional Animal Care and Use Committee (IACUC) at Columbia University Medical Center under protocol AABV0664. Mice were bred and maintained in a specific pathogen-free (SPF) facility. The Gastrin-DTR-p2A-TdTomato mouse line was generated at Columbia University, diphtheria toxin (DT) was dissolved in DPBS and administered intraperitoneally at two doses of 25 ng/20 g on alternate days to ablate antral G cells, weekly DT injections maintained hypogastrinemia.^20^ The Cck2r-CreERT2-BAC mouse line has been previously reported,^8^ and Chrm3^fl/fl^ mouse^15^ and Lgr5-DTR-eGFP mouse^51^ were described previously. LSL-hM3Dq, R26-TdTomato and R26-ZsGreen mice were obtained from Jackson Laboratory. Tamoxifen was administered by oral gavage at 2 mg/0.2 mL of corn oil per 20 g of body weight. For chronic activation of designer receptors exclusively activated by designer drugs (DREADD)-controlled neuronal activity, the ligand CNO was administered through drinking water at a concentration of 1 μg CNO per ml for 6 hours per day for 2-8 weeks. Selitrectinib (LOXO-195; 30 mg/kg/day; A285925, Ambeed Inc) was administered once daily via oral gavage. For all experiments, 8- to 12-week-old age- and sex-matched C57BL/6 mice, backcrossed for more than 10 generations, were used. For gastrin pump preparation, gastrin was dissolved in DPBS and delivered using ALZET osmotic pumps at a rate of 10 nmol/kg/hour, as previously described.^52^ Unless otherwise specified, all experiments were performed at least three times with a minimum of three biological replicates. All animals were included in the analysis unless they failed to recover post-treatment or had incomplete data due to technical issues.

#### Mouse surgery

Mice were randomly assigned to experimental and control groups to minimize bias. Randomization was performed using a random number generator to allocate animals to treatment groups. In cases where randomization was not possible, efforts were made to match groups based on age and sex. Blinding was implemented at multiple stages to minimize bias. All surgical procedures were performed under isoflurane inhalation anesthesia (2-3%), with postoperative analgesia provided by subcutaneous buprenorphine (0.1 mg/kg).

For unilateral vagotomy, a midline incision was made to access the abdominal cavity, and the anterior truncal vagus nerve was transected, resulting in selective vagal denervation of the anterior stomach while preserving pyloric function.^18^ In the control group, only the anterior vagus nerve was explored without further intervention.

Darifenacin hydrobromide was dissolved in a 50% DMSO/50% phosphate-buffered saline solution and delivered via subcutaneously implanted ALZET osmotic pumps at 6.0 mg/kg/day. Control animals received pumps filled with the solvent solution alone.^53^ At the end of the study, animals were euthanized, and the stomach was excised, opened along the greater curvature with microdissection scissors, washed with cold PBS, and pinned flat. Macroscopic images were captured, and tissues were collected for further analysis.

Fast Blue Retrograde tracing. A laparotomy was performed to expose the stomach. The greater curvature of the stomach was held with microsurgical forceps. Fast Blue (1%, Polysciences) was injected slowly at several spots beneath the serosa of the whole anterior wall or posterior wall of antrum with a Hamilton syringe (30 G, 25 μl). The needle was kept in place for 10 s to avoid dye leakage. The injection site was rinsed with saline to remove any leaking dye. Mice were euthanized 6 days after injection. The JNC was dissected.

JNG intraganglionic CCK-SAP injection. Mice were anesthetized and a midline ventral neck incision was made. The hypoglossal nerve was gently separated from the vagus nerve to expose the right jugular–nodose ganglion (JNG) swelling. 0.6 µg CCK-SAP (Advanced Targeting Systems, KIT-31) or Control-SAP was microinjected into the right JNG swelling. The incision was closed with sterile sutures. Mice were euthanized 1 week after injection, and the JNG and medulla oblongata were collected for downstream analyses.

### METHOD DETAILS

#### Ethics approval

This study involves human participants and was approved by the ethics committee of the Fujian Medical University Union Hospital (no. 2024KY233). Participants gave informed consent to participate in the study before taking part. All animal experiments and procedures were conducted following the NIH guidelines for animal research and were reviewed and approved by the Institutional Animal Care and Use Committee (IACUC) at Columbia University Medical Center under protocol AABV0664.

#### Chronic *H. pylori* infection model and MNU-induced injury model

Mice were infected with *H. pylori* strain SS1 by oral gavage of 0.2 mL brucella broth containing a cumulative dose of 100 million colony-forming units per mouse, administered across three doses per week. Successful infection was confirmed using real-time PCR. N-nitroso-N-methylurea was dissolved in distilled water at 240 ppm and freshly prepared three times per week. The solution was provided in light-shielded bottles and administered ad libitum, alternating weekly for 10 weeks, totaling 5 weeks of exposure.

#### H&E staining

For H&E staining, tissue sections were incubated in hematoxylin for 3–5 minutes, clarified for 30 seconds to 1 minute, followed by a 1-minute incubation in Bluing Reagent, and then stained with Eosin Y for 20–40 seconds.

#### Immunofluorescence

Tissues were either fixed with formalin overnight, followed by dehydration with 70% ethanol and paraffin embedding, or fixed with 4% paraformaldehyde overnight, followed by dehydration with 30% sucrose and embedding with optimal cutting temperature (O.C.T.) compound. Sections of 5 μm thickness were stained with the specified antibodies using standard protocols. Formalin-fixed, paraffin-embedded tissue sections were deparaffinized with xylene and rehydrated with phosphate-buffered saline (PBS), followed by antigen retrieval by boiling the samples in Antigen Unmasking Solution (pH 6 [H-3300] or pH 9 [H-3301]; Vector Laboratories) for 30 min.

For immunofluorescence (IF) studies, the sections were treated with blocking buffer (SP-5035, Vector Laboratories) for 30 min, incubated with primary antibodies overnight at 4 °C, and washed with PBS. The sections were then incubated with Alexa Fluor Plus 488/594/647-conjugated secondary antibodies (Thermo Fisher Scientific) for 1 h at room temperature. The sections were then mounted with an antifade mounting medium containing 4’,6-diamidino-2-phenylindole (DAPI; 0100-20, SouthernBiotech), and fluorescence was examined using an inverted fluorescence microscope BZ-X800 (Keyence).

For each mouse stomach, 4–6 paraffin or frozen sections were collected, and 6–10 randomly selected high-power fields (HPFs) per section were analyzed. The mean value of all HPFs was calculated for each individual mouse and used for group comparisons.

#### Histological evaluation

The scoring system used for histological analysis follows the methodology outlined in Cancer Res, 2005, 65 (23), pp. 10709-10715. Briefly, Histopathological evaluation was performed using H&E-stained slides, and the following criteria was used^28^: 1. Hyperplasia: Defined by increased epithelial and glandular cell density within the corpus mucosa, with epithelial cells maintaining normal morphology. 2. Metaplasia: Marked by the replacement of normal corpus glandular epithelium with cells resembling goblet cells. 3. Dysplasia: Characterized by enlarged nuclei, irregular nuclear contours, an elevated nuclear-to-cytoplasmic ratio, and atypical mitotic figures in the corpus glands. Histopathological scoring was performed by two experienced pathologists in a blind manner.

#### Gastrin Quantification

Blood samples were collected from the inferior vena cava of mice and centrifuged at 1000 g for 15 minutes at 4°C. The resulting supernatant was used for gastrin quantification. Plasma gastrin levels were measured using the Gastrin EIA Kit (Sigma-Aldrich, RAB0200) according to the manufacturer’s instructions.

#### Enzyme-linked immunosorbent assay (ELISA)

Acetylcholine (ACh) levels in mouse antral tissue were measured by ELISA. Briefly, stomachs were rapidly harvested and placed in ice-cold PBS containing the acetylcholinesterase inhibitor neostigmine (100 µM; HY-B0423, MedChem Express). After washing, the antrum was dissected, and equal tissue weights were transferred to pre-chilled microcentrifuge tubes containing four stainless-steel beads and 200 µL ice-cold RIPA buffer supplemented with neostigmine. Tissues were homogenized, centrifuged at 4°C (5,000 × g, 5 min), and clarified supernatants were collected. ACh concentrations in the supernatants were quantified using an ACh ELISA Kit (E-EL-0081, Elabscience) according to the manufacturer’s instructions.

#### Antrum dissection and single-cell isolation

Mice were euthanized by CO₂ inhalation followed by cervical dislocation. Stomachs were opened along the greater curvature and rinsed with ice-cold PBS containing 1% penicillin/streptomycin until clear. The antral mucosa was carefully dissected from the flattened stomach under a surgical microscope, cut into ∼0.5-cm pieces, and incubated in 10 mM EDTA in DPBS on ice for 60 min. Tissue fragments were then vigorously pipetted in cold Advanced DMEM/F12 supplemented with 10% FBS to release glands, and the gland-containing supernatant was collected and centrifuged (250 × g, 5 min, 4°C). The pellet was resuspended in 2 mL DPBS, supplemented with 5 mL TrypLE and 50 µL DNase I (100 mg/mL), and incubated at 37°C for 7 min with vertexing (5 s) every 2 min. Digestion was quenched by adding 2 mL CM-GF (Advanced DMEM/F12 supplemented with GlutaMAX, HEPES, and Pen/Strep) containing 10% FBS. Cells were pelleted (350 × g, 5 min, 4°C), resuspended in 3 mL CM-GF with 10% FBS, triturated, filtered through a 40-µm strainer, and centrifuged again (350 × g, 5 min, 4°C).

#### Flow cytometry and cell sorting

All FACS analyses were on LSRII or LSRFortessa instruments. Cell sorting was performed on a BD Influx cell sorter. Quantitative results were expressed as the percentage of viable EpCAM+ cells, following a standard gating strategy for epithelial cells (Cells/Single Cells/DAPI^-^EpCAM^+^) or fibroblasts (Cells/Single Cells/DAPI^-^EpCAM^-^CD31^-^PDGFa^+^).

#### Fibroblast cells culture

PDGFa⁺ fibroblasts were isolated by FACS and plated in 6-well plates. Cells were maintained in Dulbecco’s modified Eagle’s medium (DMEM) supplemented with 10% fetal bovine serum (FBS), 1% penicillin–streptomycin, 1% GlutaMAX, and basic FGF (bFGF; 25 ng/mL) at 37°C in a humidified incubator with 5% CO₂. Where indicated, cultures were treated with carbachol (10 μM) and/or darifenacin hydrobromide (2 μM). After 48 h, cells were harvested for qPCR analysis.

#### 3D Culture

Single cells were sorted by FACS based on reporter gene expression and suspended in cold Matrigel (Corning) at a density of 1000 cells per 20 μL. The cell suspension was seeded as droplets in the center of wells in a 48-well plate. The plate was incubated at 37°C for 10 minutes to allow the Matrigel to solidify. Then, the cells were overlaid with 500 μL of 50% WRN (Wnt3A, R-spondin1, and noggin) conditioned medium^54^, supplemented with 1% penicillin/streptomycin, 10 mM HEPES, 1×GlutaMAX, 1× N_2_, 1×B27, 1 mM N-acetylcysteine, and 50 ng/mL EGF, and incubated at 37°C with 5% CO_2_ to support organoid formation. For the first 3 days, 10 μM Y-27632 (ROCK inhibitor) was included. The culture media were refreshed every 1–3 days. The designated groups received 10 μM Carbachol, 2 μM Darifenacin hydrobromide, 100 nM Trametinib, 0.5 μM Verteporfin, or 100 nM gastrin, each added to the medium every other day. The area of organoids per well was quantified using microscopic imaging and BZ-X800 analyzer.

#### RNA extraction and qRT-PCR

Single sorted cells or corpus tissues from mice were lysed in RLT buffer and stored at -80°C until RNA isolation was performed, following the manufacturer’s instructions (Qiagen, 74034). cDNA synthesis was carried out using the qScript cDNA SuperMix System (QuantaBio, 95048), optimized for qRT-PCR analyses. qRT-PCR assays were performed using the SYBR Green method on the 7300 Real-Time PCR System to assess gene expression levels.

Gene expression data were normalized using the ΔΔCt method, and the results were presented as average fold changes relative to *Gapdh* expression. A comprehensive list of primer sequences is available in Table S2.

### QUANTIFICATION AND STATISTICAL ANALYSIS

#### Mouse Single Cell RNA-seq Analysis

Single Cck2r⁺ cells (ZsGreen^+^) from Cck2r-CreERT2; Rosa26-ZsGreen; Gastrin-DTR-p2A-tdTomato mice under three conditions: control (PBS), G-cell ablation (DT), and G-cell ablation with 2400ppm MNU gavage exposure (DT+MNU) were sorted by FACS 48 hours post tamoxifen induction. Immediately after sorting, cells were counted using an automated cell counter (ThermoFisher Countess II FL) to check viability and processed for library preparation (10X chromium). Library preparation and sequencing were performed by the JP Sulzberger Columbia Genome Centre (Single cell analysis core), using standard methodologies.

The Seurat R package (version 5.1.0) was used for the processing with analysis pipelines similar to those described in the human scRNA seq. quality control (QC) using the following criteria: nGene > 200 and < 8000, nUMI > 500 and < 50,000, and mitochondrial genome < 20%. After QC, samples from identical conditions underwent integration via Harmony to mitigate batch effects. We obtained 10690 cells (Ctrl, n=3011; DT, n=5353; DT+MNU, n=2326) and classified them into 7 clusters according to the top 50 marker genes (Table S1): These included a stem/progenitor cluster (*Mki67, Ccna2, Top2a, Birc5, Stmn1*); a stress-activated progenitor-like cluster enriched for p53 and stress-response genes such as *Gdf15*, *Trp53inp1*, and *Ccng1*, along with *Notch3* and *Mybl1*, which are associated with progenitor maintenance during regeneration; pre-pit and pit cell clusters (Tff1, Tff2, Muc5ac); basal gland mucous cells (*Muc6, Lgr5, Aqp5, Glp1r, Car9*); a tuft cell cluster (*Dclk1, Pou2f3*); and two endocrine clusters—endocrine 1 (*Neurod1, Gast*) and endocrine 2 (*Chga, Sstr2*).

Monocle3 trajectory analysis predicted that the progenitors serve as the origin of all other cell types, including chief cells and SPEMs. CytoTRACE has been used with default parameters for the inference of cellular differentiation states. We used Seurat’s FindMarkers function to identify differentially expressed genes (DEGs) from our scRNA-seq data. We then performed Gene Set Enrichment Analysis (GSEA) on these DEGs using the hallmark gene sets (’mh.all.v2023.2.Mm.symbols.gmt’) obtained from the Molecular Signatures Database (MSigDB) to explore the associated biological processes.

#### Human Single Cell RNA-seq Analysis

We re-analyzed single-cell RNA sequencing data from thirteen gastric antral mucosa biopsies obtained from nine patients with Non-atrophic gastritis, Chronic atrophic gastritis, Intestinal metaplasia, or Early gastric cancer (GEO: GSE134520).^30^ The Seurat R package (version 5.1.0)^44^ was used to import and process the gene expression matrices. Initial preprocessing thresholds included genes detected in at least three cells and cells with at least 300 genes. Quality filtered using the following criteria: nGene > 500 and < 6000, nUMI > 500 and < 40,000, and mitochondrial genome < 15%. The normalized data function was used to normalize the data. Principal component analysis (PCA) was performed on the top 2000 highly variable genes, and clustering analysis was performed on the top 20 PCA. Marker genes were identified using the COSG package (version 0.9.0) ^45^ and then manually annotated based on differentially expressed genes (DEGs) references from previous literature and CellMarker 2.0.^46^ The atlas mainly comprised epithelial cells (EPCAM, KRT18, and MUC1) and non-epithelial cells (VIM and PTPRC). ^30^ We further defined the epithelial cells into eight distinct cell-type clusters: antral basal gland mucous cells (GMCs, marked as MUC6 and TFF2), pit mucous cells (PMCs, marked as MUC5AC and TFF1), enteroendocrine cells (EE, CHGA and CHGB), proliferating cells (PC) representing isthmus cells (MKI67 and TOP2A), gastric cancer cells (GC, marked as CEACAM5 and CEACAM6), goblet cells (MUC2), enterocyte as a mature intestinal metaplastic cell (FABP1), and metaplastic stem-like cells (MSC) (OLFM4, EPHB2, and SOX9). CytoTRACE ^49^ has been used with default parameters for the inference of cellular differentiation states. We then performed Gene Set Enrichment Analysis (GSEA) on these DEGs using the hallmark gene sets (’h.all.v7.5.symbols.gmt’) obtained from the Molecular Signatures Database (MSigDB) to explore the associated biological processes.

#### Processing of spatial transcriptomics data

We re-analyzed spatial transcriptomics data from a gastric antral cancer tissue section obtained at Fujian Medical University Union Hospital (Genome Sequence Archive database, accession number: HRA011663).^42^ The specimen was collected from a 39-year-old male who underwent laparoscopic-assisted radical distal gastrectomy; the tumor was moderately-to-poorly differentiated with a mixed histological subtype and staged as pT3N1M0. Briefly, the tissue section was prepared and deparaffinized following the standard 10× Genomics protocol. Subsequently, a human whole-transcriptome probe set was applied to the tissue. The hybridized probes were used to construct a spatial transcriptomics library, which was then sequenced on an Illumina NovaSeq 6000 platform (service provided by Novogene Co., Ltd.). Raw sequencing data in FASTQ format, along with matched histological images, were analyzed using the 10× Genomics Space Ranger software (v1.3.0). Specifically, the FFPE “Count” pipeline with short-read probe alignment was employed to map reads to the human reference genome (GRCh38). Subsequent data processing, including gene expression normalization, dimensionality reduction, spot clustering, and identification of differentially expressed genes, was carried out using the Seurat package (v5.1.0).

#### Statistical Analysis

Statistical analyses were performed using IBM SPSS version 27.0. Data are represented as mean ± SD. The Student’s t-test was used for comparisons of two groups, and one-way ANOVA statistical evaluation was used for comparisons of at least three groups (n ≥ 3). All statistical tests were two-sided, and statistical significance was defined as p < 0.05. Experiments were repeated ≥ three times.

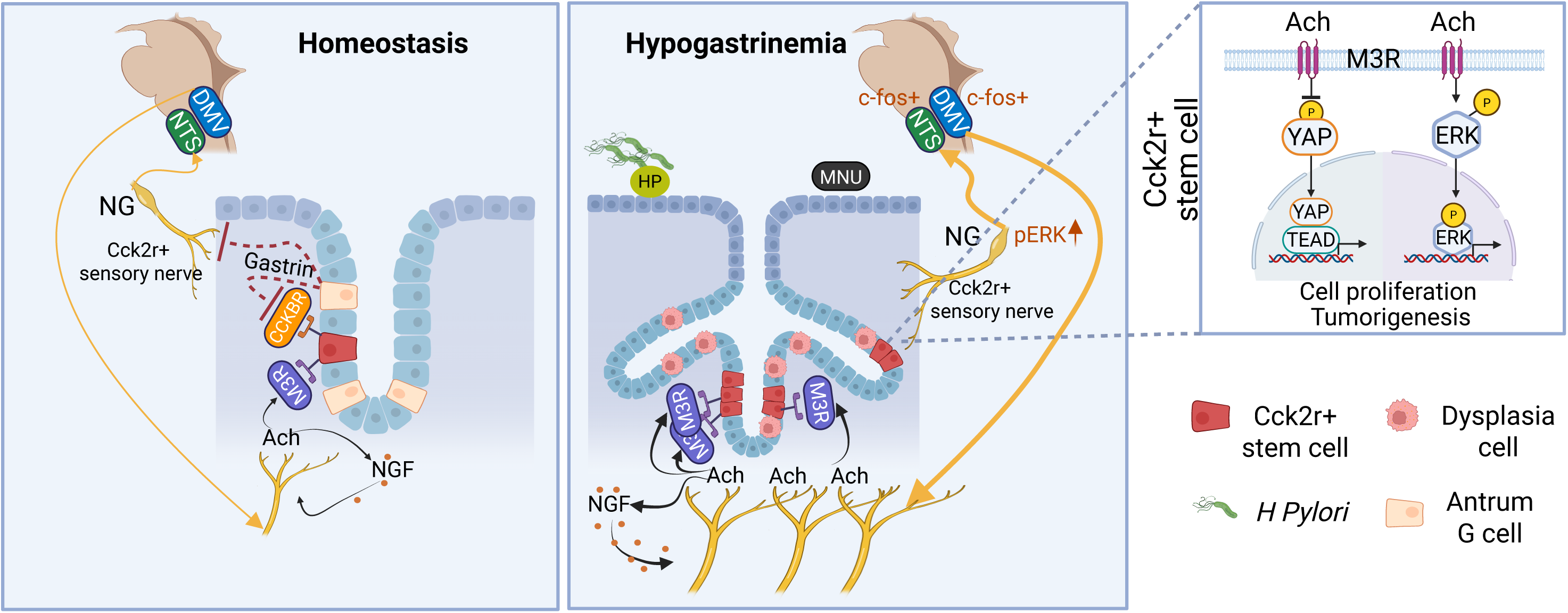

