## Supplementary material for "Endocrine–Neural Interactions Regulate Antral CCK2R⁺ Stem Cells in Gastric Inflammation and Preneoplasia": Figure S1-S7, Table S1-S2

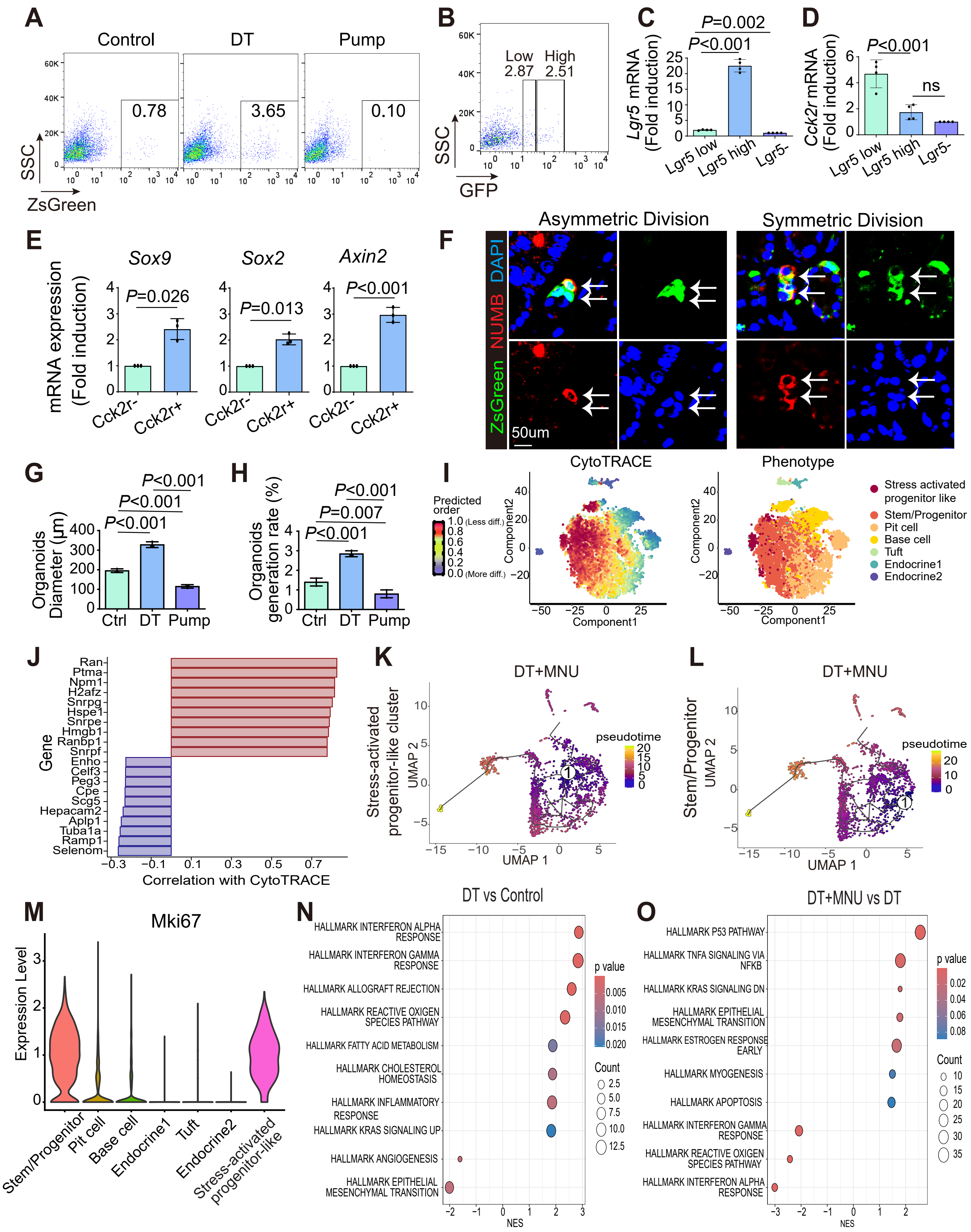
A
Control
DT
Pump
0.78
3.65
0.10
SSC
ZsGreen
B
Low 2.87
High 2.51
SSC
GFP
C
P=0.002
P<0.001
Lgr5 mRNA (Fold induction)
Lgr5 low
Lgr5 high
Lgr5-
D
P<0.001
ns
Cck2r mRNA (Fold induction)
Lgr5 low
Lgr5 high
Lgr5-
E
Sox9
Sox2
Axin2
P=0.026
P=0.013
P<0.001
mRNA expression (Fold induction)
Cck2r-
Cck2r+
F
Asymmetric Division
Symmetric Division
ZsGreen NUMB DAPI
50um
G
P<0.001
P<0.001
P<0.001
Organoids Diameter (µm)
Ctrl
DT
Pump
H
P<0.001
P=0.007
P<0.001
Organoids generation rate (%)
Ctrl
DT
Pump
I
CytoTRACE
Phenotype
Predicted order
1.0 (Less diff.)
0.0 (More diff.)
Component1
Component2
Stress activated progenitor like
Stem/Progenitor
Pit cell
Base cell
Tuft
Endocrine1
Endocrine2
J
Gene
Ran
Ptma
Npm1
H2afz
Snrpg
Hspe1
Snrpe
Hmgb1
Ranbp1
Snrpf
Enho
Celf3
Peg3
Cpe
Scg5
Hepacam2
Aplp1
Tuba1a
Ramp1
Selenom
Correlation with CytoTRACE
K
DT+MNU
Stress-activated progenitor-like cluster
pseudotime
UMAP 1
UMAP 2
L
DT+MNU
Stem/Progenitor
pseudotime
UMAP 1
UMAP 2
M
Mki67
Expression Level
Stem/Progenitor
Pit cell
Base cell
Endocrine1
Tuft
Endocrine2
Stress-activated progenitor-like
N
DT vs Control
HALLMARK INTERFERON ALPHA RESPONSE
HALLMARK INTERFERON GAMMA RESPONSE
HALLMARK ALLOGRAFT REJECTION
HALLMARK REACTIVE OXIGEN SPECIES PATHWAY
HALLMARK FATTY ACID METABOLISM
HALLMARK CHOLESTEROL HOMEOSTASIS
HALLMARK INFLAMMATORY RESPONSE
HALLMARK KRAS SIGNALING UP
HALLMARK ANGIOGENESIS
HALLMARK EPITHELIAL MESENCHYMAL TRANSITION
p value
Count
NES
O
DT+MNU vs DT
HALLMARK P53 PATHWAY
HALLMARK TNFA SIGNALING VIA NFKB
HALLMARK KRAS SIGNALING DN
HALLMARK EPITHELIAL MESENCHYMAL TRANSITION
HALLMARK ESTROGEN RESPONSE EARLY
HALLMARK MYOGENESIS
HALLMARK APOPTOSIS
HALLMARK INTERFERON GAMMA RESPONSE
HALLMARK REACTIVE OXIGEN SPECIES PATHWAY
HALLMARK INTERFERON ALPHA RESPONSE
p value
Count
NES

### Figure S2

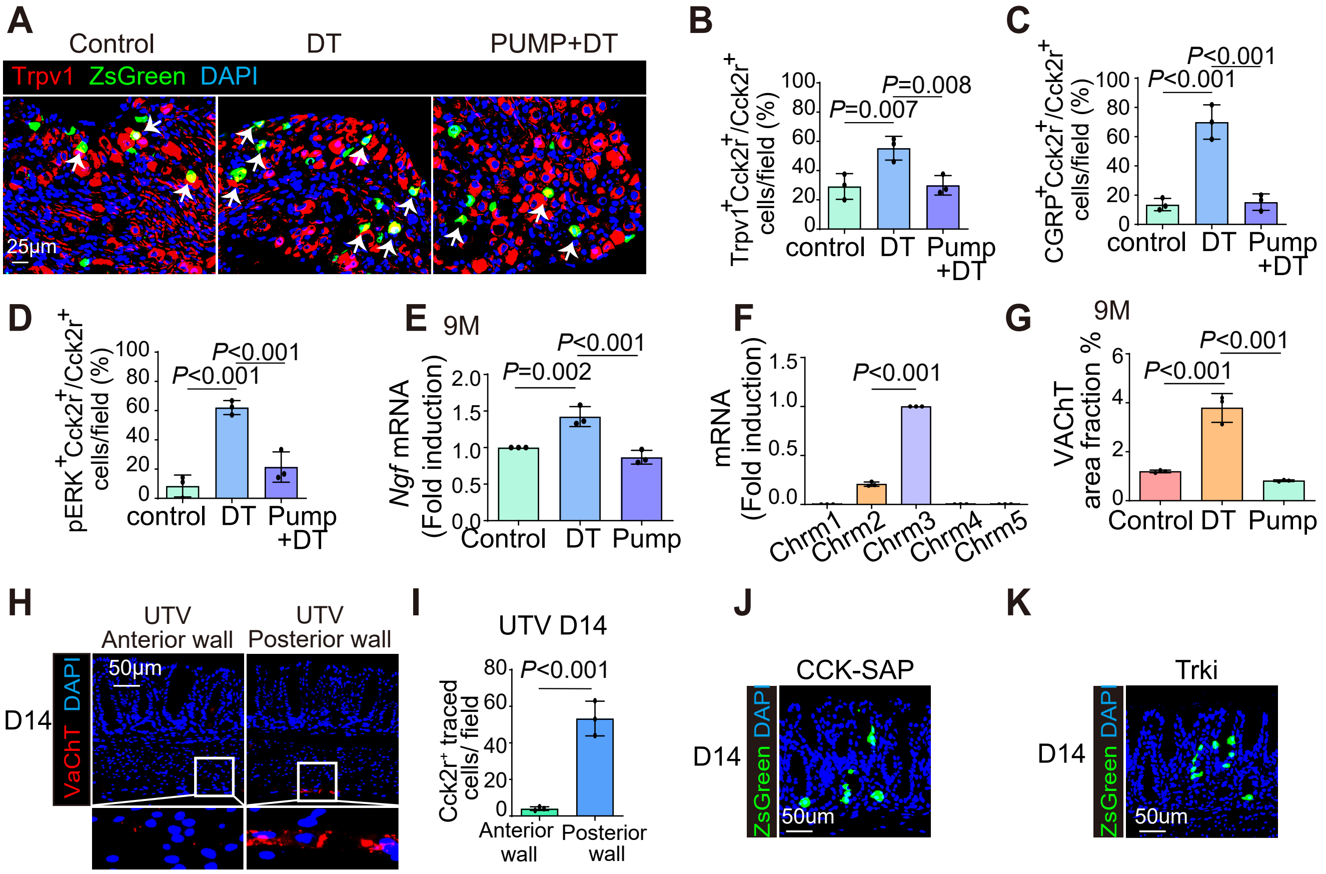
A
Control
DT
PUMP+DT
Trpv1 ZsGreen DAPI
25μm
B
Trpv1+Cck2r+/Cck2r+ cells/field (%)
P=0.007
P=0.008
control DT Pump +DT
C
CGRP+Cck2r+/Cck2r+ cells/field (%)
P<0.001
P<0.001
control DT Pump +DT
D
pERK+Cck2r+/Cck2r+ cells/field (%)
P<0.001
P<0.001
control DT Pump +DT
E
9M
Ngf mRNA (Fold induction)
P=0.002
P<0.001
Control DT Pump
F
mRNA (Fold induction)
P<0.001
Chrm1 Chrm2 Chrm3 Chrm4 Chrm5
G
9M
VAChT area fraction %
P<0.001
P<0.001
Control DT Pump
H
UTV Anterior wall
UTV Posterior wall
D14
VaChT DAPI
50μm
I
UTV D14
Cck2r+ traced cells/ field
P<0.001
Anterior wall
Posterior wall
J
CCK-SAP
D14
ZsGreen DAPI
50um
K
Trki
D14
ZsGreen DAPI
50um

### Figure S3

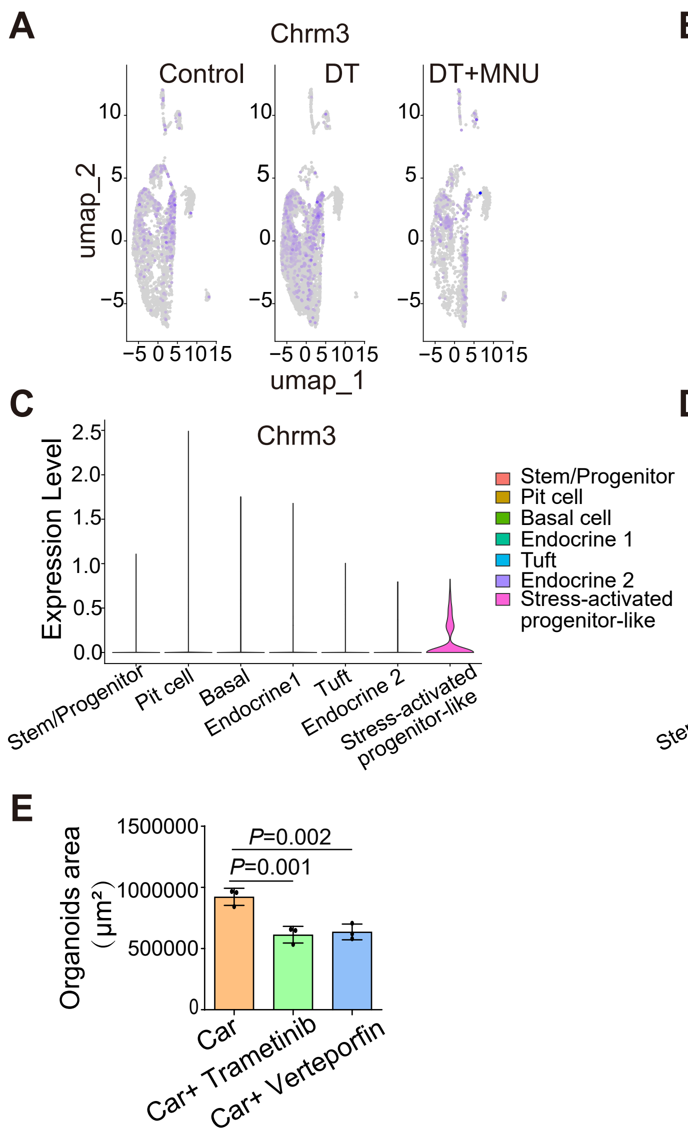

A
Chrm3
Control
DT
DT+MNU
umap_2
umap_1
B
Chrm1
Control
DT
DT+MNU
umap_2
umap_1
C
Chrm3
Expression Level
Stem/Progenitor
Pit cell
Basal cell
Endocrine 1
Tuft
Endocrine 2
Stress-activated progenitor-like
Stem/Progenitor
Pit cell
Basal
Endocrine1
Tuft
Endocrine 2
Stress-activated progenitor-like
D
Chrm1
Expression Level
Stem/Progenitor
Pit cell
Basal cell
Endocrine 1
Tuft
Endocrine 2
Stress-activated progenitor-like
Stem/Progenitor
Pit cell
Basal
Endocrine1
Tuft
Endocrine 2
Stress-activated progenitor-like
E
Organoids area (μm²)
1500000
1000000
500000
0
P=0.002
P=0.001
Car
Car+ Trametinib
Car+ Verteporfin

### Figure S4

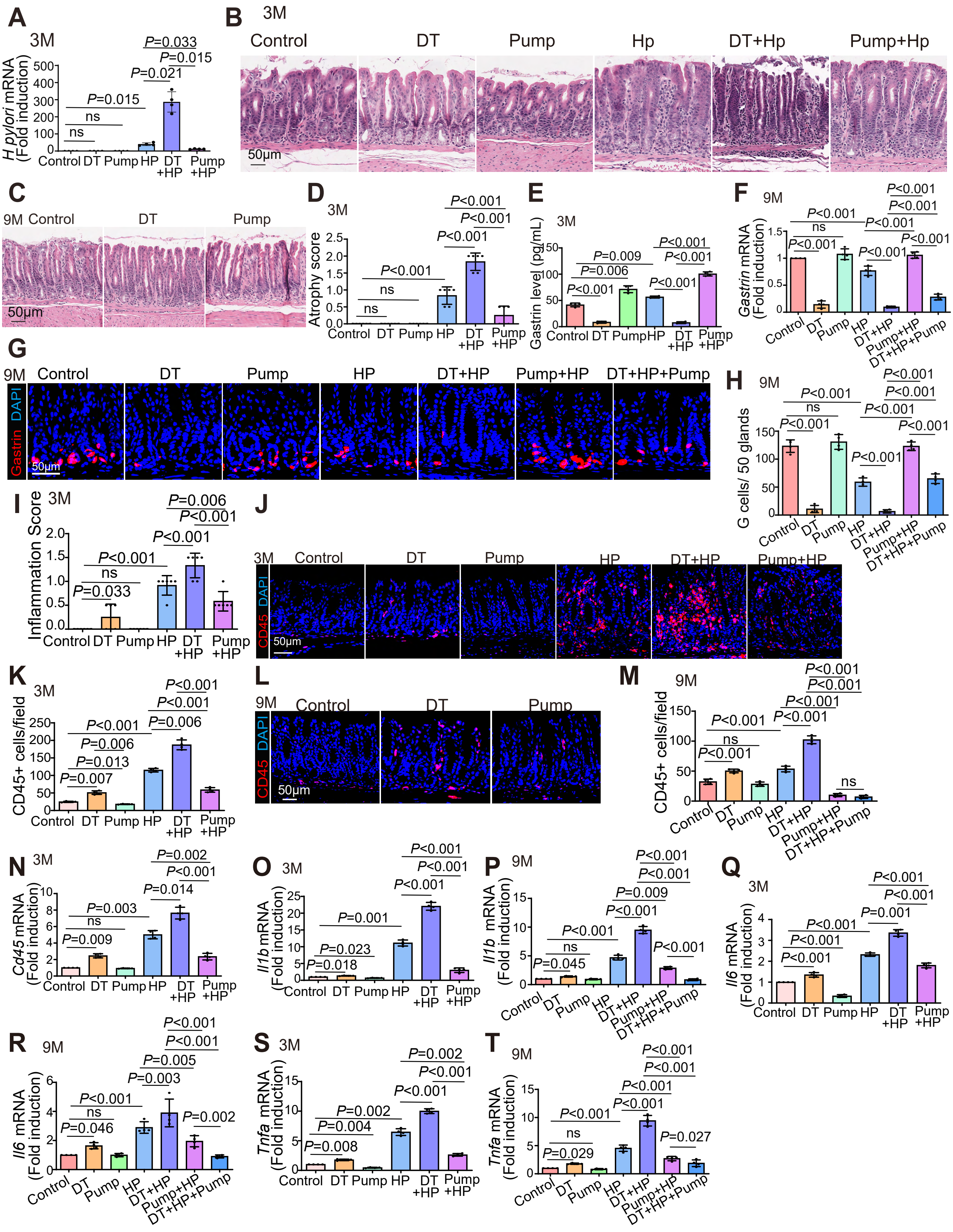
A
3M
H pylori mRNA (Fold induction)
Control DT Pump HP DT +HP Pump +HP
P=0.033
P=0.015
P=0.021
P=0.015
ns
ns
B
3M
Control
DT
Pump
Hp
DT+Hp
Pump+Hp
50μm
C
9M Control
DT
Pump
50μm
D
3M
Atrophy score
Control DT Pump HP DT +HP Pump +HP
P<0.001
E
3M
Gastrin level (pg/mL)
Control DT Pump HP DT +HP Pump +HP
P=0.009
P=0.006
F
9M
Gastrin mRNA (Fold induction)
Control DT Pump HP DT+HP Pump+HP DT+HP+Pump
G
9M
Control
DT
Pump
HP
DT+HP
Pump+HP
DT+HP+Pump
DAPI
Gastrin
50μm
H
9M
G cells/ 50 glands
I
3M
Inflammation Score
P=0.006
P=0.033
J
3M
CD45
DAPI
K
3M
CD45+ cells/field
P=0.013
P=0.007
L
9M
M
9M
N
3M
Cd45 mRNA (Fold induction)
P=0.002
P=0.014
P=0.003
P=0.009
O
3M
Il1b mRNA (Fold induction)
P=0.001
P=0.023
P=0.018
P
9M
P=0.045
Q
3M
Il6 mRNA (Fold induction)
R
9M
P=0.005
P=0.046
S
3M
Tnfa mRNA (Fold induction)
P=0.004
P=0.008
T
9M
P=0.027
P=0.029

### Figure S5

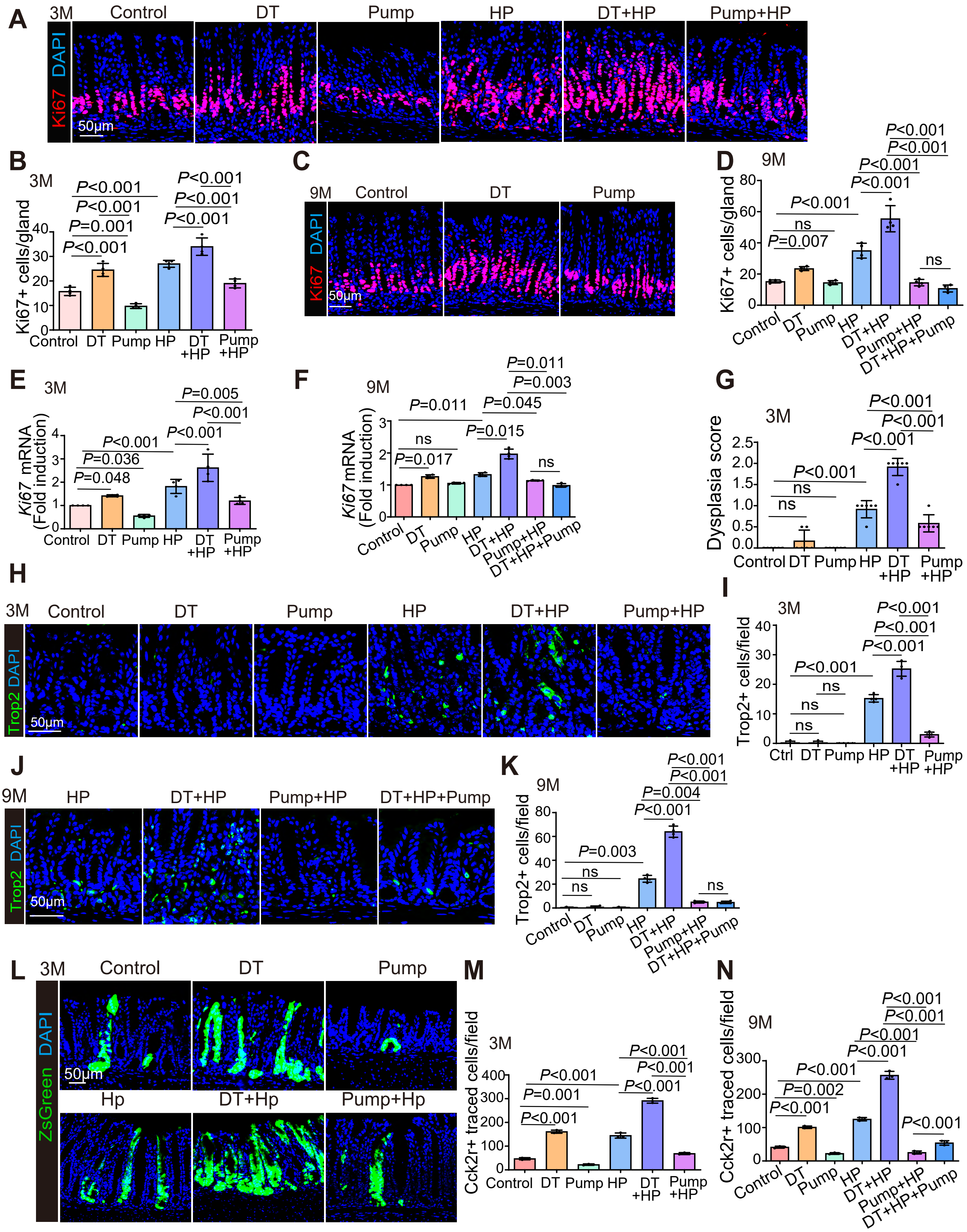

A
3M
Control
DT
Pump
HP
DT+HP
Pump+HP
Ki67 DAPI
50μm
B
C
D
E
F
G
H
I
J
K
L
M
N
Ki67+ cells/gland
Ki67 mRNA (Fold induction)
Dysplasia score
Trop2+ cells/field
Cck2r+ traced cells/field
Trop2 DAPI
ZsGreen DAPI
9M
DT+HP+Pump
Ctrl
Hp
DT+Hp
Pump+Hp

### Figure S6

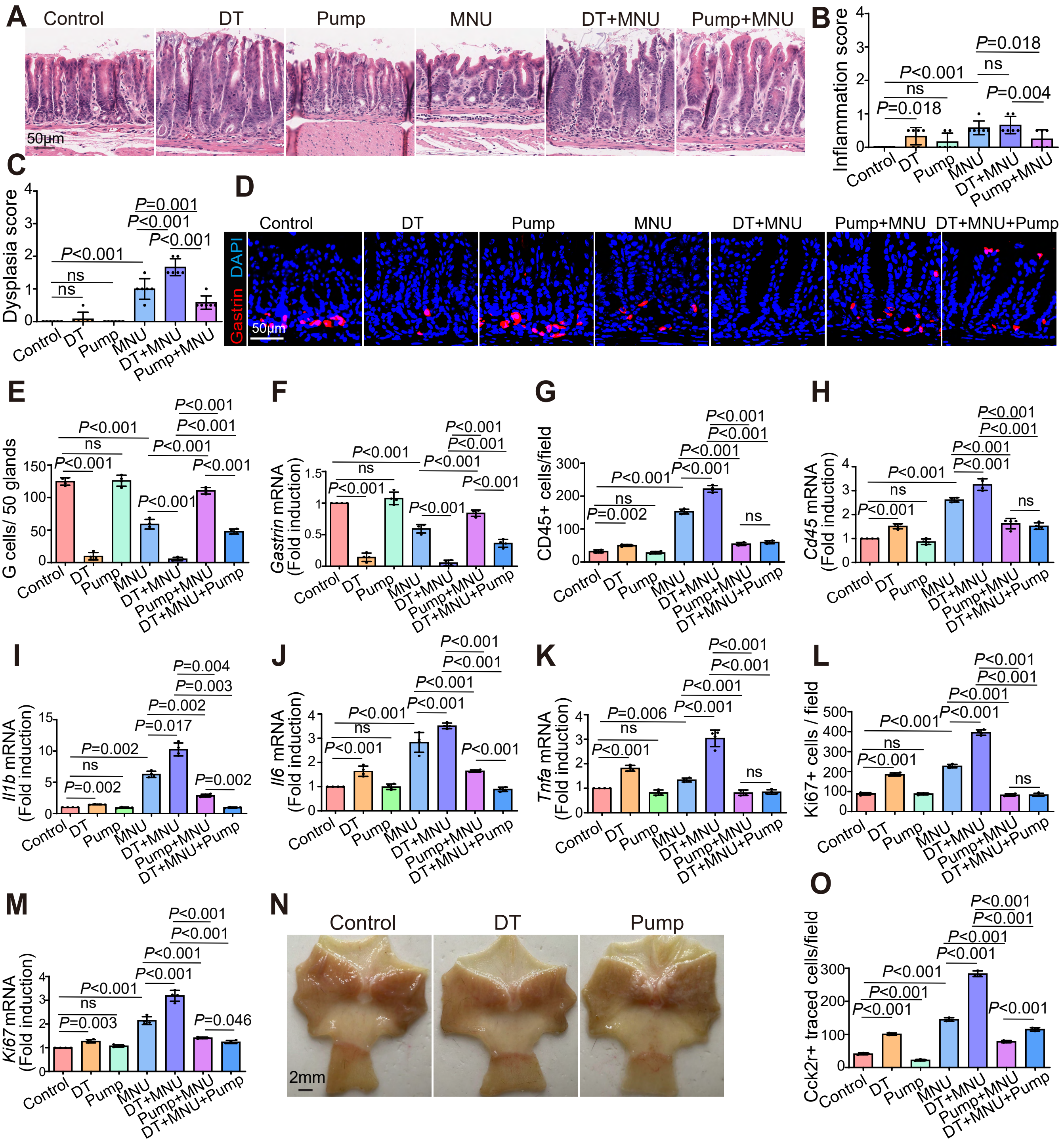

A
Control
DT
Pump
MNU
DT+MNU
Pump+MNU
50μm
B
Inflammation score
C
Dysplasia score
D
Gastrin DAPI
DT+MNU+Pump
E
G cells/ 50 glands
F
Gastrin mRNA (Fold induction)
G
CD45+ cells/field
H
Cd45 mRNA (Fold induction)
I
Il1b mRNA (Fold induction)
J
Il6 mRNA (Fold induction)
K
Tnfa mRNA (Fold induction)
L
Ki67+ cells / field
M
Ki67 mRNA (Fold induction)
N
2mm
O
Cck2r+ traced cells/field

#### Figure S7

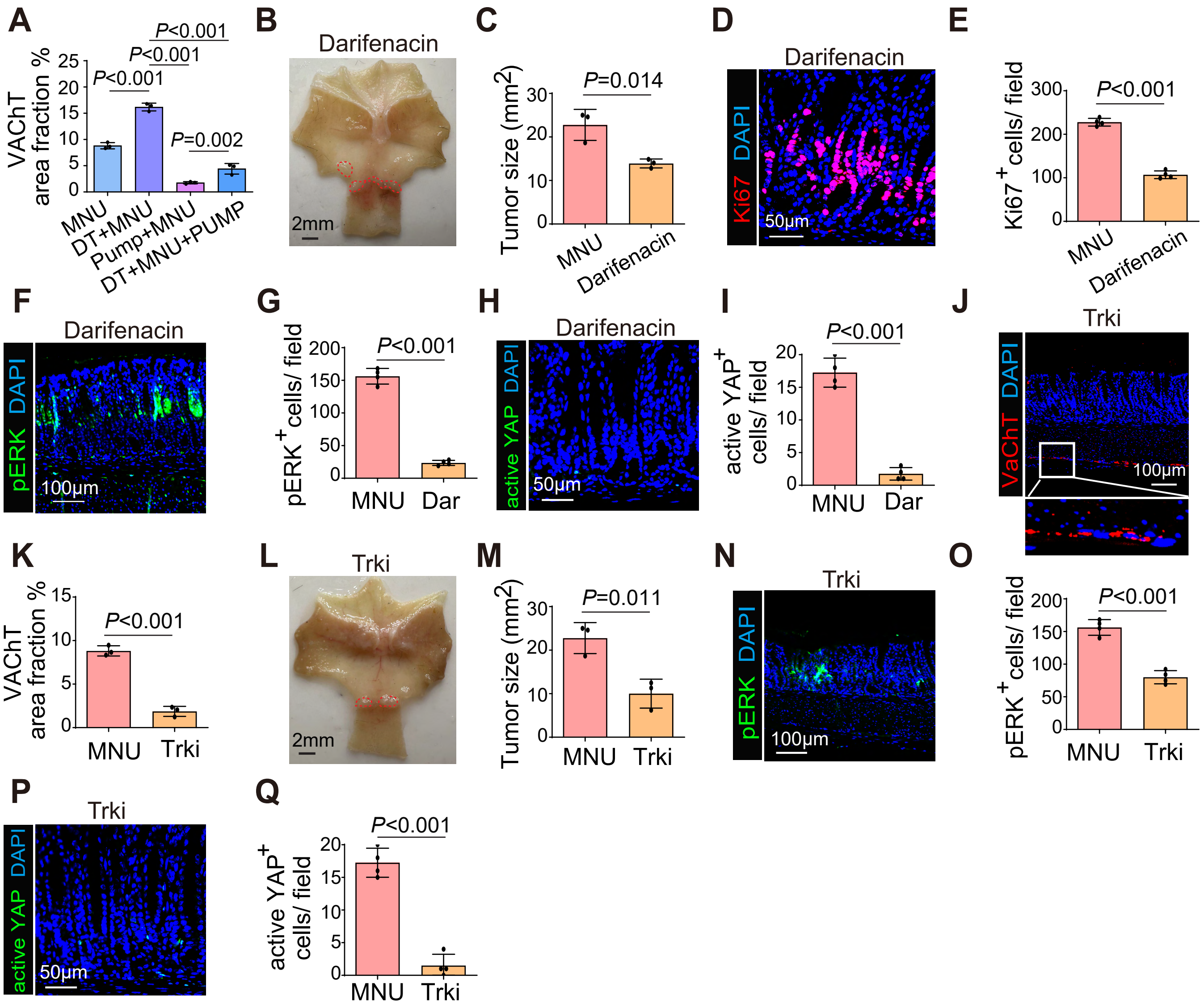

A
VAChT area fraction %
P<0.001
P<0.001
P<0.001
P=0.002
MNU
DT+MNU
Pump+MNU
DT+MNU+PUMP
B
Darifenacin
2mm
C
Tumor size (mm2)
P=0.014
MNU
Darifenacin
D
Darifenacin
Ki67 DAPI
50μm
E
Ki67+cells/ field
P<0.001
MNU
Darifenacin
F
Darifenacin
pERK DAPI
100μm
G
pERK+cells/ field
P<0.001
MNU
Dar
H
Darifenacin
active YAP DAPI
50μm
I
active YAP+ cells/ field
P<0.001
MNU
Dar
J
Trki
VaChT DAPI
100μm
K
VAChT area fraction %
P<0.001
MNU
Trki
L
Trki
2mm
M
Tumor size (mm2)
P=0.011
MNU
Trki
N
Trki
pERK DAPI
100μm
O
pERK+cells/ field
P<0.001
MNU
Trki
P
Trki
active YAP DAPI
50μm
Q
active YAP+ cells/ field
P<0.001
MNU
Trki

#### Table S1. Top 50 marker genes for each cluster in mouse antral scRNA-seq.

| | Stem/progenitor 1 | Pit | Basal cell | Pre-pit cell | Basal cell | Stress-activated progenitor-like | Tuft | Endocrine 1 | Endocrine 2 |
|---|---|---|---|---|---|---|---|---|---|
| | Cluster 1 | Cluster 3 | Cluster 4 | Cluster 3 | Cluster 4 | Cluster 2 | Cluster 5 | Cluster 6 | Cluster 7 |
| | names.0 | names.1 | names.2 | names.3 | names.4 | names.5 | names.6 | names.7 | names.8 |
| 1 | Pclaf | Sptssb | Aqp5 | AY036118 | Pdia2 | Gdf15 | Lrmp | Pcsk1n | Ptf1a |
| 2 | Pbk | Aldh3a1 | A4gnt | Tff1 | Pla2g1b | Cox6b2 | Rgs13 | Neurod1 | Pnliprp1 |
| 3 | Smc2 | Gm3776 | Slc9a3 | Gkn2 | Glp1r | Eda2r | Fyb | Pax6 | Vtn |
| 4 | Mki67 | Mucl3 | Muc6 | Ptma | Ppp1r3a | AU040972 | Rgs2 | Runx1t1 | Hdc |
| 5 | Lmnb1 | Gsta1 | Creb3l4 | Gkn1 | Kcnj15 | Phlda3 | Gnat3 | Cryba2 | Lhfpl5 |
| 6 | H2afx | Lgals3 | Copz2 | Tpt1 | Pga5 | Ddit4l | Dclk1 | Vwa5b2 | Slc18a2 |
| 7 | Hmgb2 | 2210407C18Rik | Wfdc18 | Tff2 | Npw | Inka2 | Gng13 | Scgn | Rbpjl |
| 8 | Tmpo | Smim24 | Lrg1 | H2afz | Esrrg | Ano3 | Hck | Nkx2-2 | Gc |
| 9 | Tk1 | Mal | Rgs5 | Tmsb4x | Lypd6 | 1700007K13Rik | Sh2d6 | Syt13 | Cel |
| 10 | Ccna2 | Rep15 | 2010007H06Rik | Eef1a1 | Gpm6a | Abcb1b | Alox5ap | Fam183b | Kcnk3 |
| 11 | Tuba1b | Vstm2b | Acot1 | S100a6 | Car9 | Ces2e | 1810046K07Rik | Isl1 | Kcnmb2 |
| 12 | Rrm1 | Slc5a5 | Snhg18 | H3f3b | Pdzd2 | Gask1b | Pik3r5 | Map1b | Chga |
| 13 | Cks1b | Akr1b8 | Chad | Ppia | Scgb2b7 | Gria3 | Pik3cg | Dbpht2 | Nupr1 |
| 14 | Cks2 | Anxa3 | Lox | Agr2 | Kcne2 | 9230114K14Rik | Spib | Insm1 | Prune2 |
| 15 | Top2a | Gldc | Rsph9 | Gkn3 | 2210404E10Rik | Tubb6 | Hpgds | Fev | P2rx1 |
| 16 | Racgap1 | Adh7 | F5 | Gsta4 | Basp1 | Zfp365 | Sh2d7 | Cpe | Car4 |
| 17 | Lig1 | Gsta2 | Col11a2 | Actg1 | Tuba3a | Trp53inp1 | Alox5 | Smarca1 | Plvap |
| 18 | Asf1b | Lypd8 | Cldn2 | Hspa8 | Arhgap24 | Slc25a48 | Pou2f3 | Tubb3 | Pappa2 |
| 19 | Smc4 | Mllt3 | Rnf152 | Krt8 | Tinag | Tomm20l | Ly6g6f | Scg3 | Hspb1 |
| 20 | Cdca8 | S100a14 | Slc12a8 | Naca | Ak4 | Cd80 | Trpm5 | Hap1 | Rab3c |
| 21 | Cdk1 | Syt8 | Mup10 | Fau | Igf1 | Anxa8 | Ptgs1 | Gng4 | Kirrel2 |
| 22 | Dek | Gm3336 | Muc3a | Gm42418 | Mogat1 | Psrc1 | Nrep | Gast | Insrr |
| 23 | Aurkb | Oasl1 | Sord | Txn1 | Arhgef4 | Ddias | Hmx2 | Cldn4 | Ncam1 |
| 24 | Gmnn | Muc5ac | 4931406C07Rik | Prdx1 | Dnah8 | Ccng1 | Strip2 | Sphkap | Col11a1 |

| | | | | | | | | | |
|---|---|---|---|---|---|---|---|---|---|
| 25 | Kif11 | Capg | Padi2 | Fth1 | P3h4 | Tnfsf4 | Rac2 | Cd177 | Cckar |
| 26 | Mcm7 | Lgals4 | Mup14 | Eef1b2 | Nr4a2 | Kank3 | Bmx | Scin | Matn4 |
| 27 | Birc5 | Gsn | Cblif | S100a11 | Smim31 | Exoc4 | Vav1 | Pcsk1 | Rgs7bp |
| 28 | Dut | Ahnak | Furin | Rack1 | Arhgdig | Tnfrsf18 | Hmx3 | A2ml1 | Ccn3 |
| 29 | Rrm2 | Phgr1 | Pigr | Hmgb2 | Synm | Mgmt | Matk | Cacna1a | Sez6 |
| 30 | Hist1h2ae | Sqor | Spdef | Chchd2 | Nr2f2 | Tex15 | Kctd12 | Nefm | Fam89a |
| 31 | Lsm2 | Sdcbp2 | Vldlr | Dbi | 4930404N11Rik | Mcam | Pate4 | Kcnb2 | Prkcb |
| 32 | Stmn1 | Psca | Serpina3n | Actb | Usp53 | Sesn2 | Ltc4s | Serpini1 | Gm11837 |
| 33 | Hist1h2ap | Sult1b1 | C2cd4b | Tmsb10 | Ppp2r2b | 9530053A07Rik | Pygl | Pnmal1 | Slc39a5 |
| 34 | Knl1 | 2200002D01Rik | Cd44 | Tubb5 | Tmem151a | Notch3 | Adh1 | Baiap3 | Camk1d |
| 35 | Lsm3 | AA467197 | C4b | Slc25a5 | Slc1a3 | Scn1b | Nrgn | Stxbp5l | 1810010K12Rik |
| 36 | Melk | Gsdma2 | Tpd52l1 | Cystm1 | Ltbp4 | Gm42047 | Cd300lf | Cldn6 | Nrp1 |
| 37 | Kif23 | Ethe1 | Gm830 | Cdk1 | Slc9b2 | S100a3 | Zfp428 | Rfx6 | Pnmal2 |
| 38 | Rfc5 | Ces2c | Car12 | Hint1 | Bpifb1 | Acaa1b | Tspan6 | Pyy | Cpb2 |
| 39 | Shcbp1 | Il18 | Itln1 | Car2 | Pnliprp2 | Ggta1 | Rgs22 | Gm14964 | Tnr |
| 40 | Tpx2 | Gkn2 | Tspan12 | Eif1 | 1010001N08Rik | Mdm2 | Plcb2 | Gm609 | Gm27199 |
| 41 | Ska1 | Sytl2 | Lgr5 | Ubb | Ppp3ca | Gna15 | Inpp5d | Spock2 | Sstr2 |
| 42 | Esco2 | Ptgr1 | Hgfac | Cox5a | Fam3c | Mybl1 | Espn | Serping1 | Stmn3 |
| 43 | Spc25 | Sult1d1 | Habp2 | Btf3 | 1700016K19Rik | Atg9b | A4galt | Emb | Cckbr |
| 44 | Hist1h1b | Capn9 | Slc12a2 | Sprr2a3 | Cwh43 | Glipr1 | Mctp1 | Ptprn | Reg1 |
| 45 | Gins2 | Cdc42ep2 | Aqp1 | Gpx2 | Fgfbp1 | Ccp110 | Siglecf | Acsl6 | Slc38a3 |
| 46 | Tubb5 | Gkn1 | Etv5 | Ran | Aldh6a1 | Zmat3 | Ptpn6 | Gnao1 | Wnt4 |
| 47 | Pcna | Mpst | Pla2g5 | H3f3a | Emp1 | C330021F23Rik | Inpp5j | Arx | Hamp2 |
| 48 | Nasp | Krt19 | Itih2 | Tagln2 | Atp7b | Ckap2 | Plcg2 | Ttyh1 | Neurl1a |
| 49 | Tyms | Bcas1 | Slc16a7 | Hspe1 | Cnn3 | Pvt1 | Limd2 | Gpx3 | Esr1 |
| 50 | Gemin6 | Slurp1 | Tmem45a | Ftl1 | Dtna | Zfp385a | Gprc5c | Ret | Enho |

**Table S2. Sequences of qRT-PCR Primers, related to STAR Methods.**

| Gene | Forward primer (5'-3') | Reverse primer (5'-3') |
|---|---|---|
| ***Gapdh*** | AGGTCGGTGTGAACGGATTTG | TGTAGACCATGTAGTTGAGGTCA |
| ***H.pylori SS1*** | CAAAATCGCTGGCATTGGT | CTTCACCGGCTAAGGCTTCA |
| ***Ngf*** | TGATCGGCGTACAGGCAGA | GCTGAAGTTTAGTCCAGTGGG |
| ***Lgr5*** | CTCCACACTTCGGACTCAACAG | AACCAAGCTAAATGCACCGAAT |
| ***Cck2r*** | GCCGTTTCCTACCTCATGGGGGT | CCCTTGGTTTCGGACCCGGC |
| ***Sox9*** | GTGCAAGCTGGCAAAGTTGA | TGCTCAGTTCACCGATGTCC |
| ***Sox2*** | GCGGAGTGGAAACTTTTGTCC | CGGGAAGCGTGTACTTATCCTT |
| ***Axin2*** | ATAAGCAGCCGTTCGCGATG | CAGCAATCGGCTTGGTCTCTC |
| ***Gastrin*** | ACACAACAGCCAACTATTC | CAAAGTCCATCCATCCGTAG |
| ***Cd45*** | TTTCCAATGTGCTGTGTCCT | TGAAGAAGAGAGATCCACCCA |
| ***Chrm1*** | CAGAAGTGGTGATCAAGATGCCTAT | GAGCTTTTGGGAGGCTGCTT |
| ***Chrm2*** | TGGAGCACAACAAGATCCAGAAT | CCCCTGAACGCAGTTTTCA |
| ***Chrm3*** | CCAGTTGGTGTGTTCTTCCTT | AGGAAGAGCTGATGTTGGGA |
| ***Chrm4*** | GTGACTGCCATCGAGATCGTAC | CAAACTTTCGGGCCACATTG |
| ***Chrm5*** | GGCCCAGAGAGAACGGAAC | TTCCCGTTGTTGAGGTGCTT |
| ***Ki67*** | ATCATTGACCGCTCCTTTAGGT | GCTCGCCTTGATGGTTCCT |
| ***Il-1β*** | GCAACTGTTCCTGAACTCAACT | ATCTTTTGGGGTCCGTCAACT |
| ***Il-6*** | TAGTCCTTCCTACCCCAATTTCC | TTGGTCCTTAGCCACTCCTTC |
| ***Tnf-a*** | CAGGCGGTGCCTATGTCTC | CGATCACCCCGAAGTTCAGTAG |

**Supplementary Figure Legends**

**Supplementary Figure 1. Flow-cytometric validation and extended single-cell analyses support stemness and stress-associated reprogramming of antral Cck2r$^+$ cells.** (A) Representative flow cytometric image of Cck2r$^+$ epithelial cells in the antrum one day after tamoxifen induction. (B) Representative flow cytometric image of Lgr5$^{hi}$ and Lgr5$^{low}$ epithelial cells in the antrum of Lgr5DTR-eGFP mouse. (C, D) qRT-PCR quantification of *Lgr5* (C) and *Cck2r* (D) mRNA. (E) qRT-PCR quantification of *Sox9, Sox2, Axin2* mRNA. (F) Representative images of asymmetric versus symmetric division in paired ZsGreen$^+$ cells two days post-tamoxifen, Scale bars: 50 μm. (G-H) Diameter (G), and formation efficiency (H) of organoids from FACS-sorted Cck2r$^+$ cells after 7 days of culture. (I) CytoTRACE heatmap of transcriptionally defined clusters, with higher scores indicating less differentiated, more stem-like states. (J) Bar plot showing top genes positively (red) and negatively (blue) correlated with CytoTRACE scores. (K, L) Pseudotime trajectory analysis of scRNA-seq data, showing developmental transitions from undifferentiated (blue-violet) to differentiated (yellow) states from stress activated progenitor-like cluster (K) and stem/progenitor cluster (L). (M) Violin plot of *MKi67* expression across Cck2r$^+$ cell clusters. (N, O) Gene set enrichment analysis (GSEA) of hallmark pathways in epithelial cells, comparing DT versus control (N) and DT+MNU versus DT (O). Data represent mean ± SD ($n \geq 3$). DT, diphtheria toxin; MNU, N-nitroso-N-methylurea; Pump, continuous gastrin infusion via osmotic pump.

**Supplementary Figure 2. Gastrin loss induces vagal cholinergic remodeling and NGF-dependent activation of antral CCK2R$^+$ stem cells.** (A) Co-immunofluorescence of Trpv1 (red) with ZsGreen (green) in the NG. Scale bars, 25 μm. (B–D) Quantification of Trpv1$^+$ (B), CGRP$^+$ (C), and pERK$^+$ (D) cells among ZsGreen⁺ (Cck2r-lineage–labeled) cells in the NG. (E) Relative *Ngf* mRNA expression in antral tissue determined by qPCR. (F) qRT–PCR analysis of muscarinic receptor transcripts M1–M5 (Chrm1–Chrm5) in FACS-sorted antral Pdgfra⁺ fibroblasts. (G) Quantification of VAChT⁺ cholinergic fibers in the antrum after 9 months of G-cell ablation and 3 months of gastrin pump infusion. (H, I) Representative immunofluorescence image (H) and quantification (I) of VAChT⁺ cholinergic fibers in the antrum 14 days after UTV. Scale bars, 50 μm. (J, K) Representative image of Cck2r⁺ stem-cell lineage tracing in the antrum 14 days after CCK-SAP (J) and Trki (K) treatment. Scale bars, 50 μm. All experiments were performed using Cck2r-CreERT2; Rosa26-ZsGreen; Gastrin-DTR-p2A-TdTomato mice unless otherwise indicated. Data are presented as mean ± SD (n ⩾ 3). DT, diphtheria toxin; Pump, continuous gastrin infusion via osmotic pump; UTV, unilateral truncal vagotomy.

**Supplementary Figure 3. Vagus Nerve–Muscarinic Receptor 3 Pathway Mediates Antral tumor initiation.** (A, B) UMAP feature plots of muscarinic receptor gene expression, showing selective enrichment of *Chrm3* (A) and low *Chrm1*(B) in antral Cck2r⁺ cells. (C, D) Violin plot of *Chrm3* (C) and *Chrm1* (D) expression across transcriptionally defined Cck2r⁺ cell clusters from scRNA-seq. (E) Quantification of organoid area from FACS-sorted antral Cck2r⁺ single cells cultured for 6 days under indicated treatments. Data are presented as mean ± SD (n ≥ 3 mice per group). Car, carbachol; DT, diphtheria toxin; Pump, continuous gastrin infusion; MNU, N-nitroso-N-methylurea;

**Supplementary Figure 4. G-cell ablation intensifies *H. pylori*–induced antral inflammation and atrophy.** (A) qRT–PCR quantification of *H. pylori* colonization at 3 months post-infection. (B, C) Representative H&E images of the antrum at 3 months (B) and 9 months (C) post-infection. (D) Quantification of antral atrophy scores at 3 months post-infection. (E) Plasma gastrin levels at 3 months post-infection. (F) Antral *Gastrin* mRNA expression at 9 months post-infection. (G, H) Representative gastrin immunofluorescence images (G) and quantification (H) at 9 months post-infection. (I) Histological inflammation scores in the antrum at 3 months post-infection. (J, K) $CD45^+$ immune-cell infiltration at 3 months, assessed by immunofluorescence (J) and quantification (K). (L, M) $CD45^+$ immune-cell infiltration at 9 months, assessed by immunofluorescence (L) and quantification (M). (N–T) qRT–PCR analysis of inflammatory transcripts in the antrum, including *Cd45* (N), *Il1b* at 3 months (O) and 9 months (P), *Il6* at 3 months (Q) and 9 months (R), and *Tnfa* at 3 months (S) and 9 months (T) post-infection. All experiments were performed in the antrum of Cck2r-CreERT2; Rosa26-ZsGreen; Gastrin-DTR-p2A-TdTomato mice. Scale bars, 50 μm. Data are mean ± SD ($n \geq 3$). 3M, 3 months; 9M, 9 months; DT, diphtheria toxin; Pump, continuous gastrin infusion via osmotic pump; HP, *Helicobacter pylori*.

**Supplementary Figure 5. G-cell ablation enhances *H. pylori*–induced antral proliferation and dysplasia.** (A, B) Ki67 immunofluorescence in the antrum at 3 months post-infection (A) and quantification (B). (C, D) Ki67 immunofluorescence at 9 months post-infection (C) and quantification (D). (E, F) qRT–PCR analysis of *Mki67* mRNA at 3 months (E) and 9 months (F) post-infection. (G) Histological dysplasia scores in the antrum at 3 months post-infection. (H, I) Trop2 immunofluorescence at 3 months post-infection (H) and quantification (I). (J, K) Trop2 immunofluorescence at 9 months post-infection (J) and quantification (K). (L, M) Representative images (L) and quantification (M) of $Cck2r^+$ lineage tracing in the antrum at 3 months after tamoxifen induction. (N) Quantification of $Cck2r^+$ lineage tracing at 9 months after tamoxifen induction. All experiments were performed in the antrum of Cck2r-CreERT2; Rosa26-ZsGreen; Gastrin-DTR-p2A-

TdTomato mice. Scale bars, 50 μm. Data are mean ± SD ($n \geq 3$). DT, diphtheria toxin; Pump, continuous gastrin infusion via osmotic pump; Hp, *Helicobacter pylori*.

**Supplementary Figure 6. Chronic G-cell ablation potentiates MNU-driven antral inflammation, dysplasia, and Cck2r$^+$ stem-cell hyperactivation.** (A) Representative H&E images of the antrum at 18 weeks post-MNU treatment. (B, C) Quantification of histological inflammation (B) and dysplasia (C) scores in the antrum at 18 weeks post-MNU treatment. (D, E) Gastrin immunofluorescence at 9 months post-MNU treatment (D) and quantification (E). (F) qRT–PCR quantification of *Gastrin* mRNA. (G) Quantification of CD45$^+$ immune-cell infiltration. (H–K) qRT–PCR analysis of inflammatory transcripts in the antrum at 9 months post-MNU treatment (*Cd45* (H), *Il1b* (I), *Il6* (J), and *Tnfa* (K)). (L, M) Quantification of Ki67$^+$ cells per field (L) and qRT–PCR quantification of *Mki67* mRNA (M). (N) Gross stomach images from control, DT, and Pump groups at 9 months post-MNU treatment. (O) Quantification of Cck2r$^+$ lineage tracing at 9 months post-MNU treatment. All experiments were performed in the antrum of Cck2r-CreERT2; Rosa26-ZsGreen; Gastrin-DTR-p2A-TdTomato mice. Scale bars, 50 μm (A, D) and 2 mm (N). Data are mean ± SD ($n \geq 3$). DT, diphtheria toxin; Pump, continuous gastrin infusion via osmotic pump; MNU, *N*-nitroso-*N*-methylurea.

**Supplementary Figure 7. Pharmacologic blockade of M3R or Trk signaling restrains vagal cholinergic remodeling, ERK/YAP activation, and MNU-driven antral tumor progression.** (A) Quantification of VAChT$^+$ cholinergic fibers in the antrum after 9 months of treatment. (B, C) Representative gross images (B) and quantification (C) of MNU-induced antral tumors following darifenacin treatment during the final month. (D, E) Ki67 immunofluorescence (D) and quantification (E) of epithelial proliferation in the antrum. (F, G) pERK immunofluorescence (F) and quantification

(G) in the antrum. (H, I) Active YAP immunofluorescence (H) and quantification (I) in the antrum. (J, K) VAChT immunofluorescence (J) and quantification (K) of cholinergic fibers following Trk inhibitor (Trki) treatment. (L, M) Representative gross images (L) and quantification (M) of MNU-induced antral tumors following Trki treatment during the final month. (N, O) pERK immunofluorescence (N) and quantification (O) in the antrum after Trki treatment. (P, Q) Active YAP immunofluorescence (P) and quantification (Q) in the antrum after Trki treatment. Data are mean ± SD (n ≥ 3). All experiments were performed in the antrum of Cck2r-CreERT2; Rosa26-ZsGreen; Gastrin-DTR-p2A-TdTomato mice. Scale bars: 100 μm (F, J, N) and 50 μm (D, H, P). Dar, darifenacin; DT, diphtheria toxin; MNU, *N*-nitroso-*N*-methylurea; Pump, continuous gastrin infusion; Trki, Trk inhibitor.
